# Genetic Engineering of *Treponema pallidum* subsp. *pallidum*, the Syphilis Spirochete

**DOI:** 10.1101/2021.05.07.443079

**Authors:** Emily Romeis, Lauren Tantalo, Nicole Lieberman, Quynh Phung, Alex Greninger, Lorenzo Giacani

## Abstract

**Background:** Despite more than a century of research, genetic manipulation of *Treponema pallidum* subsp. *pallidum* (*T. pallidum*), the causative agent of syphilis, has not been successful. The lack of genetic engineering tools has severely limited understanding of the mechanisms behind *T. pallidum* success as a pathogen. A recently described method for *in vitro* cultivation of *T. pallidum,* however, has made it possible to experiment with transformation and selection protocols in this pathogen. Here, we describe an approach that successfully replaced the *tprA* (*tp0009*) pseudogene in the SS14 *T. pallidum* strain with a kanamycin resistance (*kan*^R^) cassette.

**Principal findings:** A suicide vector was constructed using the pUC57 plasmid backbone. In the vector, the *kan*^R^ gene was cloned downstream of the *tp0574* gene promoter. The *tp0574*prom-*kan*^R^ cassette was then placed between two 1-kbp homology arms identical to the sequences upstream and downstream of the *tprA* pseudogene. To induce homologous recombination and integration of the *kan*^R^ cassette into the *T. pallidum* chromosome, *in vitro*-cultured SS14 strain spirochetes were exposed to the engineered vector in a CaCl_2_-based transformation buffer and let recover for 24 hours before adding kanamycin-containing selective media. Integration of the *kan*^R^ cassette was demonstrated by qualitative PCR, droplet digital PCR (ddPCR), and whole-genome sequencing (WGS) of transformed treponemes propagated *in vitro* and *in vivo*. ddPCR analysis of RNA and mass spectrometry confirmed expression of the *kan*^R^ message and protein in treponemes propagated *in vitro*. Moreover, *tprA* knockout (*tprA*^ko^-SS14) treponemes grew in kanamycin concentrations that were 64 times higher than the MIC for the wild-type SS14 (wt-SS14) strain and in infected rabbits treated with kanamycin.

**Conclusion:** We demonstrated that genetic manipulation of *T. pallidum* is attainable. This discovery will allow the application of functional genetics techniques to study syphilis pathogenesis and improve syphilis vaccine development.

**Author Summary:** Syphilis is still an endemic disease in many low- and middle-income countries, and it has been resurgent in high-income nations for almost two decades. In endemic areas, syphilis causes significant morbidity and mortality, particularly when its causative agent, the spirochete *Treponema pallidum* subsp*. pallidum* (*T. pallidum*) is transmitted to the fetus during pregnancy. A better understanding of *T. pallidum* biology and syphilis pathogenesis would help devise better control strategies for this infection. One of the limitations associated with working with *T. pallidum* was our inability to genetically alter this pathogen to evaluate the function of genes encoding virulence factors or create attenuated strains that could be useful for vaccine development. Here, we report a transformation protocol that allowed us to replace a specific region of the *T. pallidum* genome containing a pseudogene (i.e., a non-functional gene) with a stably integrated kanamycin resistance gene. To our knowledge, this is the first-ever report of a method to achieve a genetically modified *T. pallidum* strain and, as such, it can revolutionize research in the syphilis field.

## Introduction

Syphilis is a chronic sexually transmitted infection that still represents a significant burden for public health as it causes significant morbidity and mortality worldwide. The World Health Organization (WHO) estimates that syphilis global incidence ranges between 5.6 to 11 million new cases every year, while disease prevalence is between 18 to 36 million cases worldwide [1, 2]. Although most of those cases occur in low- and middle-income countries where the disease is endemic, syphilis rates have been steadily increasing in high-income countries for decades now, including in the US. In these countries, mainly men who have sex with men (MSM) and persons living with HIV (PLHIV) are affected [3–8]. In the US, the rate of early syphilis in 2019 (11.9 cases per 100,000 population), represented a 460% increase compared to the cases reported in 2000 (2.1 cases per 100,000 population) [3]. If untreated, syphilis can progress to affect the patient’s cardiovascular and central nervous systems, possibly leading to serious manifestations such as aortic aneurism, stroke, hearing or visual loss, dementia, and paralysis [9]. Because *T. pallidum* can cross the placental barrier, mother-to-child transmission of syphilis during pregnancy accounts for up to 50% of stillbirths in sub-Saharan Africa and a high proportion of perinatal morbidity and mortality cases [10].

A better understanding of *T. pallidum* biology and syphilis pathogenesis would help in devising more effective measures for disease control. Recently, Edmondson *et al.* [11] described a method to continually propagate *T. pallidum in vitro* using a cell culture-based system previously pioneered by Fieldsteel *et al.* [12]. This method represented a major advancement in the field, in that it provided investigators with an alternative to the propagation of treponemal strains in laboratory rabbits. Despite such advancement, a limitation in the study of *T. pallidum* remained the lack of tools for genetic modification of this pathogen. The availability of the cultivation system, however, paved the way to experimenting with transformation and selection procedures to introduce foreign DNA into the *T. pallidum* genome.

Here, we describe a protocol that allowed us to integrate a kanamycin resistance (*kan*^R^) cassette into a pseudogene (*tprA*, encoded by the *tp0009* gene) of the *T. pallidum* SS14 strain. In the SS14 strain, the *tprA* gene is non-functional due to a frame-shift mutation caused by a CT dinucleotide deletion at position 712 of the annotated gene open reading frame (ORF), even though syphilis strains with a functional *tpr*A gene are known, such as the Sea81-4 strain [13]. The choice to derive a *tprA* knockout (*tprA*^ko^-SS14) strain was driven by the high likelihood that a) this region would not affect *T. pallidum* viability if removed, being already non-functional in the wild-type SS14 (wt-SS14) strain, and that b) eliminating this pseudogene would not result in a polar effect inhibiting transcription of downstream genes. The *tprA* locus, even when encoding a functional gene, is likely to be transcribed as a monocistronic mRNA based on prediction software such as Operon-mapper (https://biocomputo.ibt.unam.mx/operon_mapper/). To this end, we used a pUC57-based suicide vector where the *kan*^R^ gene was placed between two ∼1 kbp homology arms identical to the regions upstream (998 bp) and downstream (999 bp) of the *tprA* frameshifted ORF, respectively. To drive expression of the *kan*^R^ gene, the promoter of the *tp0574* gene (encoding the 47 kDa lipoprotein), previously identified by Weigel *et al.* [14], was chosen based on experimental evidence that *tp0574* is among the most highly transcribed genes in *T. pallidum*, and is possibly expressed constitutively in this pathogen [15, 16]. Following transformation using a CaCl_2_-based buffer and selection, qualitative PCR was used to confirm integration of the *tp0574*prom-*kan*^R^ construct within the *tprA* locus by priming from sequences outside of the homology arms not cloned into the vector. Transformants were shown to grow *in vitro* in a kanamycin concentration (200 µg/ml) 64 times higher than the minimal inhibitory concentration (MIC) of this antibiotic for the wt-SS14 strain (3.1 µg/ml), as well as in rabbits infected intratesticularly (IT) and treated with pharmaceutical-grade kanamycin twice a day for 10 consecutive days post-infection. Replacement of the *tprA* pseudogene with the *tp0574*prom-*kan*^R^ construct was confirmed by whole-genome sequencing from *in vitro-*cultivated treponemes as well as quantitative droplet digital PCR (ddPCR) targeting the *kan*^R^ gene and the *tp0574*, *tp0001* (*dnaA*), and *tprA* genes in the treponemal chromosome. Message levels of the *tp0574* and *kan*^R^ genes, transcribed in the *tprA*^ko^-SS14 by the same promoter, were also evaluated by RT-ddPCR from *in vitro*-grown strains. Expression of the Kan^R^ protein in the *tprA*^ko^-SS14 strain but not in wt-SS14 was demonstrated by mass spectrometry (MS).

Although we ablated a pseudogene, which did not allow us to obtain a mutant lacking a known phenotype to be evaluated through functional assays, the ability to transform and manipulate *T. pallidum* using CaCl_2_ and an appropriately engineered vector is a significant step forward in the field. Our discovery opens numerous possibilities, including classical genetic studies in this pathogen, the long-awaited application of functional genomics techniques, and even the possibility of targeting virulence factors responsible for immune evasion and persistence to obtain an attenuated strain for vaccine development.

## Results

### Transformation and selection of *T. pallidum*

The pUC57-based p*tprA*arms-*tp0574*prom-*kan*^R^ plasmid construct was used to transform wt-SS14 treponemes. Transformed *T. pallidum* cells were subsequently propagated in kanamycin-supplemented media (25 µg/ml, effective on selection experiments of other spirochetes). Because the transformation buffer contained CaCl_2_ to increase membrane permeability and facilitate plasmid intake, we also exposed wt-SS14 cells to transformation buffer alone (without plasmid, to exclude CaCl_2_ lethality for the treponemes) and proceeded to propagate these cells in culture media with no antibiotic. Furthermore, to ensure that non-transformed treponemes would not survive exposure to kanamycin at the concentration used for *in vitro* selection, the wt-SS14 strain was also propagated in media containing 25 µg/ml of kanamycin. Due to the long generation time of the SS14 strain (∼44 hours), *T. pallidum* cells were sub-cultured every 14 days instead of every week until Passage #7 (Week 12 post-transformation; Fig.1), and weekly thereafter. Transformed *tprA*^ko^-SS14 treponemes could be microscopically counted 2 weeks post-transformation (Fig.1; Passage #2), even though only 2.4 treponemes per dark-field microscope (DFM) field could be seen at this time (corresponding to a concentration of 2.4×10^6^ *T. pallidum* cells/ml). For transformed treponemes, cell density increased four weeks post-transformation and remained steady throughout Passage #6 (Week 10 post-transformation). During this window (Passage #2-6), the average number of treponemes counted was 2.8×10^7^ cells/ml. The density of wt-SS14 treated with CaCl_2_ alone was higher than that of *tprA*^ko^-SS14 cells already at Passage #2 (Week 2 post-exposure; Fig.1), suggesting that treponemes were not harmed by CaCl_2_. Propagation of this strain was halted at Passage #6 (Week 10 post-exposure; Fig.1). Wild-type SS14 cells propagated in kanamycin-containing media could not be seen on the DFM over 10 weeks of propagation, confirming the ability of 25 µg/ml kanamycin to inhibit *T. pallidum* growth *in vitro*. During Passage #7-10, the *tprA*^ko^-SS14 treponemal inoculum was increased to obtain enough cells for subsequent experiments. During these passages, the average number of treponemal cells counted was 2.8×10^8^ cells/ml. Overall, these data support that a kanamycin-resistant strain was obtained due to the transformation of the wt-SS14 strain with the p*tprA*arms-*tp0574*prom-*kan*^R^ vector.

**Figure 1.**
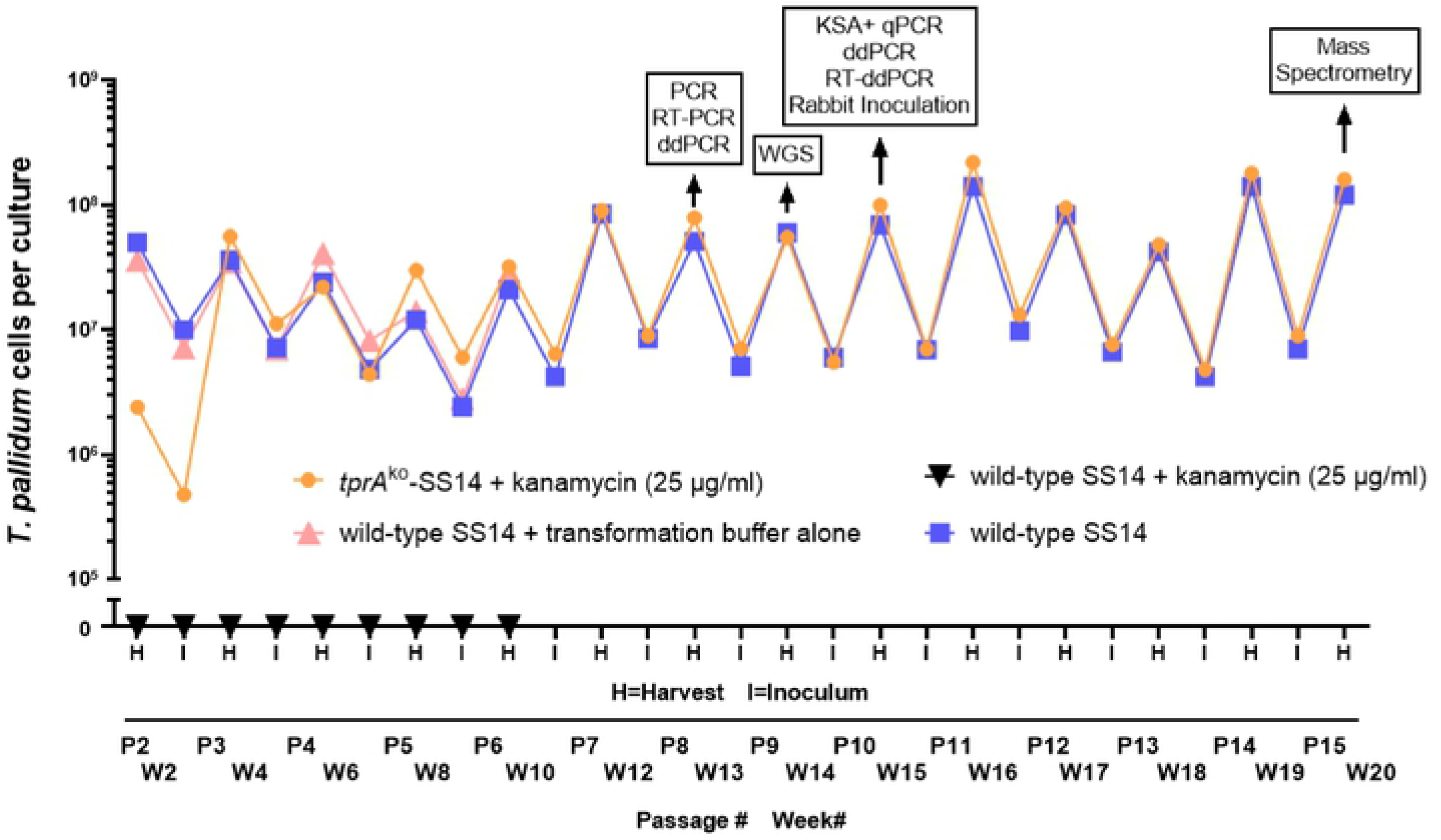
Twenty-week *in vitro* growth curve of SS14 *T. pallidum* cells post-exposure to p*tprA*arms-*tp0574*prom-*kan*^R^ plasmid in transformation buffer (orange), transformation buffer alone (pink), media containing kanamycin (black), compared to the wild-type SS14 strain (blue). RT-PCR: reverse-transcription-PCR; ddPCR: droplet digital PCR; RT-ddPCR: reverse-transcription droplet digital PCR; WGS: whole-genome sequencing; KSA: kanamycin susceptibility assay; qPCR: quantitative PCR.

### Qualitative PCR, and quantitative ddPCR to confirm integration of the *kan*^R^ gene and qualitative RT-PCR to evaluate *kan*^R^ gene expression

At Passage #8 (Week 13 post-transformation; Fig.1), *tprA*^ko^-SS14 cells were harvested and processed for a) PCR to confirm integration of the *kan*^R^ gene into the *tprA* locus, b) RT-PCR to assess the presence of message for the *kan*^R^ gene, and c) for quantitative ddPCR to evaluate the ratio between the *kan*^R^ gene and three other targets: *tprA* (*tp0009*), *dnaA* (*tp0001*), and *tp0574*. Samples from the wt-SS14 propagated in parallel to the *tprA*^ko^-SS14 strain were used as control. A qualitative, long-range PCR approach was first used to confirm integration of the *tp0574*prom-*kan*^R^ sequence into the *tprA* locus, according to the schematic reported in Fig.2A. To this end, primers annealing to the *T. pallidum* genomic region flanking the *tprA* homology arms of the construct and to the *kan*^R^ gene, respectively, were employed. In these reactions, DNA samples extracted from *tprA*^ko^-SS14 cells as well as from wt-SS14 were amplified with primer pairs 1+2, 3+4, 1+3, 5+6, and 2+4 (Fig.2A; primer sequences in Table 1). Amplification of the *tp0574* gene was performed as a positive control. As expected, amplification using primer pairs 1+2 and 3+4 (Fig.2B, sub-panels a and b, respectively) yielded a positive result only when the *tprA*^ko^-SS14 DNA was used as the template, showing that integration of the *kan*^R^ gene occurred. Amplification of the *kan*^R^ gene was positive only from DNA extracted from the *tprA*^ko^-SS14, and the transformation plasmid DNA, used as positive control (sub-panel c), while negative amplification using the 5+6 primer pair (annealing to the vector backbone; sub-panel d) showed no residual plasmid in the *tprA*^ko^-SS14 culture and that the *kan*^R^ amplicon (in sub-panel c) was not due to residual vector used for transformation weeks earlier. When used together, primers 2+4 generated a 3,746 bp amplicon with the *tprA*^ko^-SS14 strain DNA template (sub-panel e), which was the expected size if the small *kan*^R^ gene (816 bp in size) replaced the *tprA* pseudogene (1,821 bp), and a 4,643 bp amplicon in the wild-type and undetectable residual wt-SS14 in the *tprA*^ko^-SS14 culture wells. As expected, the amplification of the *tp0574* gene was uniformly positive for both the transformed and wild-type strain (sub-panel f). Overall, these data showed that the *kan*^R^ gene was integrated into the *tprA*^ko^-SS14 strain genome in place of the *tprA* pseudogene and no residual plasmid could be amplified from the culture.

**Figure 2.**
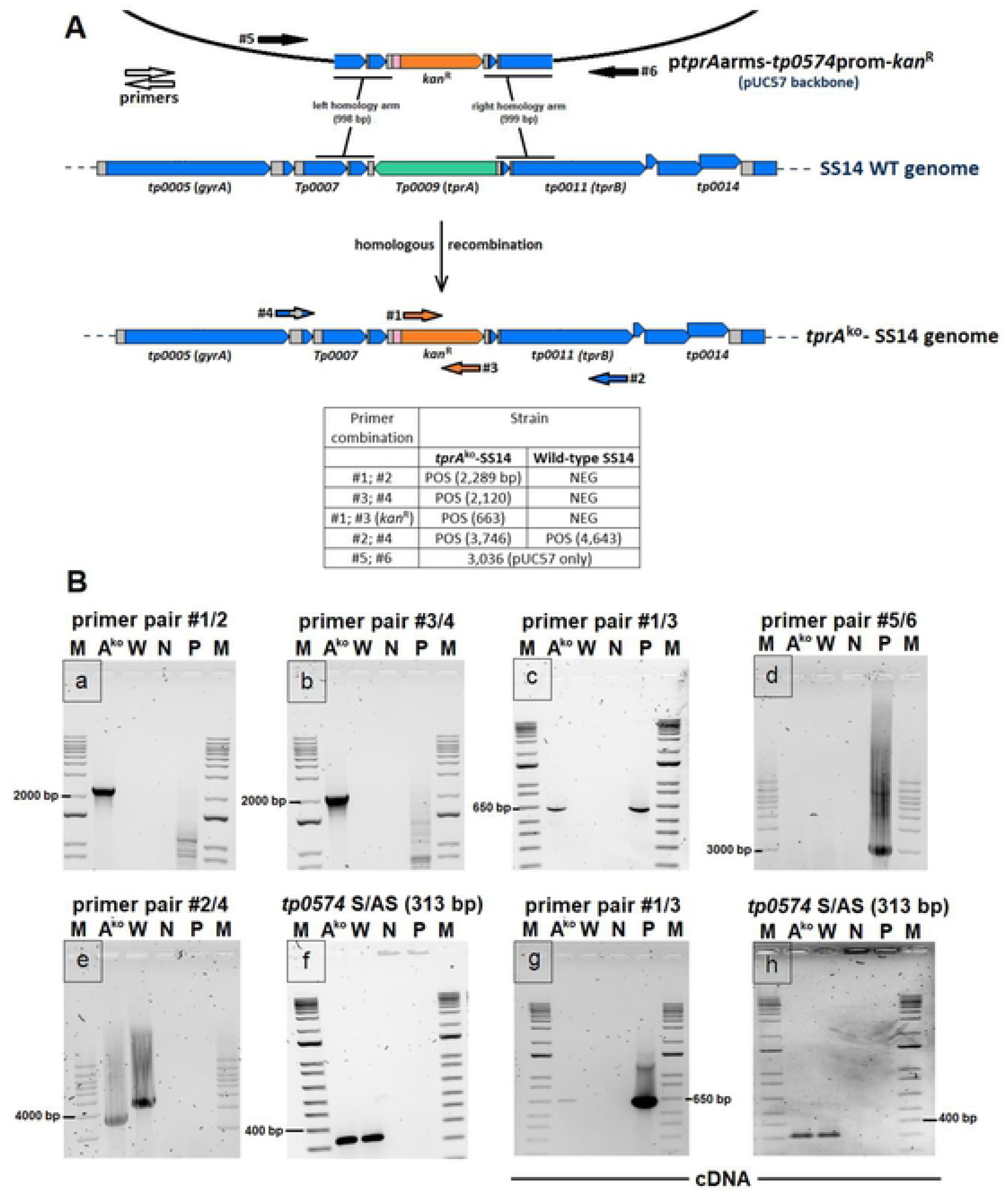
(**A**) Schematic of the recombination event that led to the *tprA*^ko^-SS14 strain. Primer positions are indicated by color-coded arrows. Amplicon sizes (bp) generated by the different primer combinations are reported in the legend. (**B**) Amplification reactions using the primer combinations in panel A on DNA (sub-panels a-f) or cDNA (sub-panels g and h) template from the *tprA*^ko^- and wt-SS14 strain harvested at Passage #8 post-transformation. M: molecular size marker (bp); A^ko^: *tprA*^ko^-SS14, W: wt-SS14; N: no-template control, P: p*tprA*arms-*tp0574*prom-*kan*^R^ plasmid DNA template.

**Table 1.**
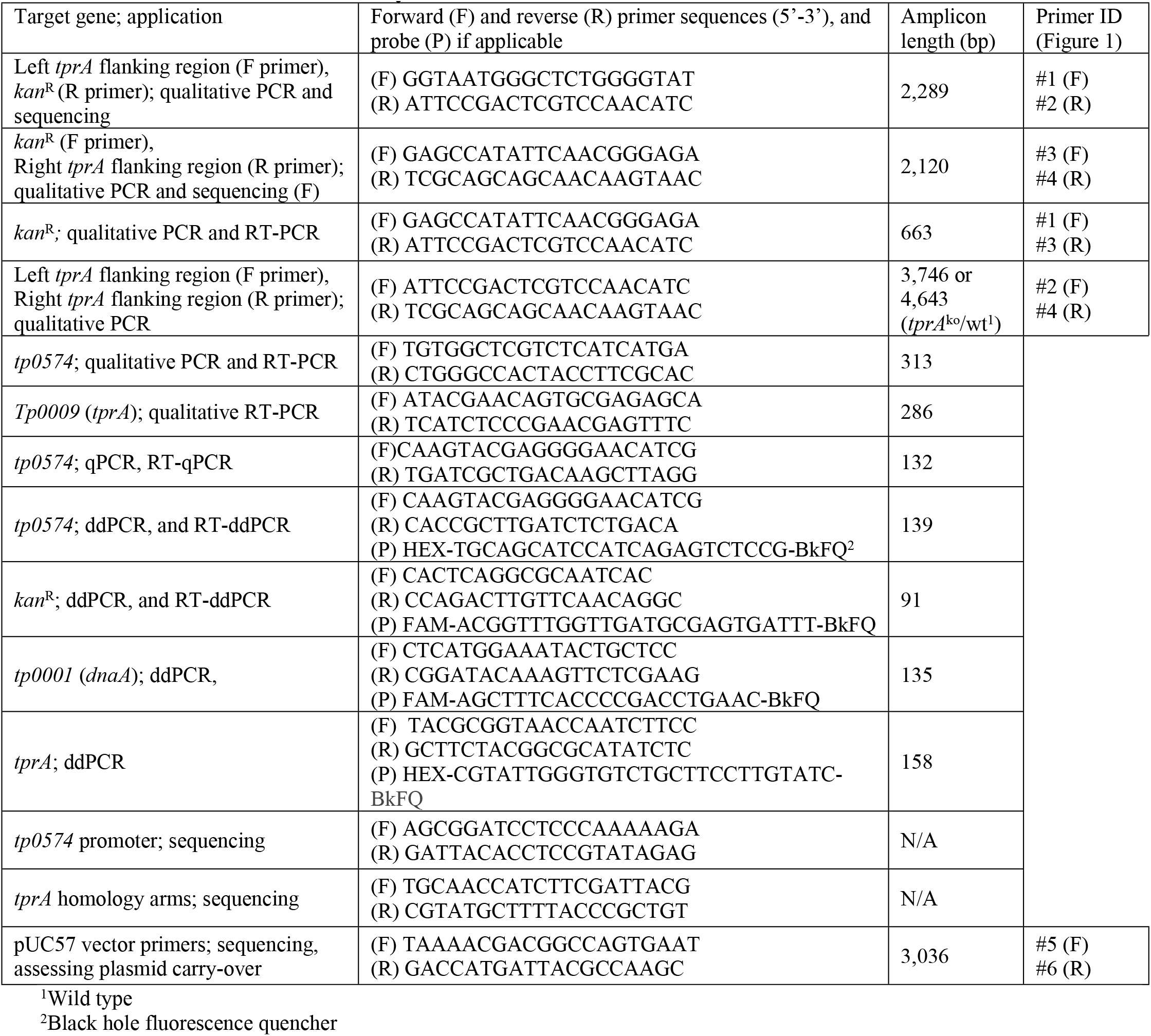
Primers used in this study

RNA extracted from the *tprA*^ko^-SS14 and wt-SS14 strain (Passage 8, Week 13 post-transformation; Fig.1) was DNaseI-treated to eliminate residual DNA and reverse transcribed. cDNA was used as template to assess transcription of the *kan*^R^ gene and *tp0574* genes, as well as the *tprA* gene. Previous studies on other *T. pallidum* strains with a *tprA* pseudogene suggested that this locus is transcribed at a very low level in these strains, even though the coding sequence contains a frameshift that would truncate the resulting peptide during translation [13]. Results (Fig.2B) showed that the *kan*^R^ gene is expressed only in the *tprA*^ko^-SS14 (sub-panel g). As expected, *tp0574* was expressed in both strains (sub-panel h). *tprA*-specific mRNA could not be detected in either sample from the *tprA*^ko^- and wt-cultures harvested at this time point; however, in samples harvested in subsequent passages, *tprA*-specific message was detected in wt-SS14 propagated alongside the *tprA*^ko^-SS14 strain, but never in the *tprA*^ko^-SS14 strain. These results supported that the *kan*^R^ transgene is actively transcribed from the *tp0574* promoter in the *tprA*^ko^-SS14 strain while the *tprA* pseudogene is no longer transcribed in this strain due to *tprA* ablation.

We next performed droplet digital PCR (ddPCR) on *kan^R^, dnaA,* and *tprA* loci in a separate laboratory to evaluate copy number ratios among these genes. In the *tprA*^ko^-SS14 strain, the *kan*^R^:*dnaA* ratio was equal to 1.05, while the *tprA*:*dnaA* ratio was virtually zero (0.006; Fig.3). These data reiterated that a) integration of the *kan*^R^ gene occurred in the *tprA* locus, b) that this replacement was stable, and c) that no extra copies of the *kan*^R^ gene existed outside of the *T. pallidum* genome. On the contrary, when the wt-SS14 DNA was used as template, the *tprA*:*dnaA* was 1.01, while the *kan*^R^:*dnaA* ratio was also virtually zero (0.005; Fig.3).

**Figure 3.**
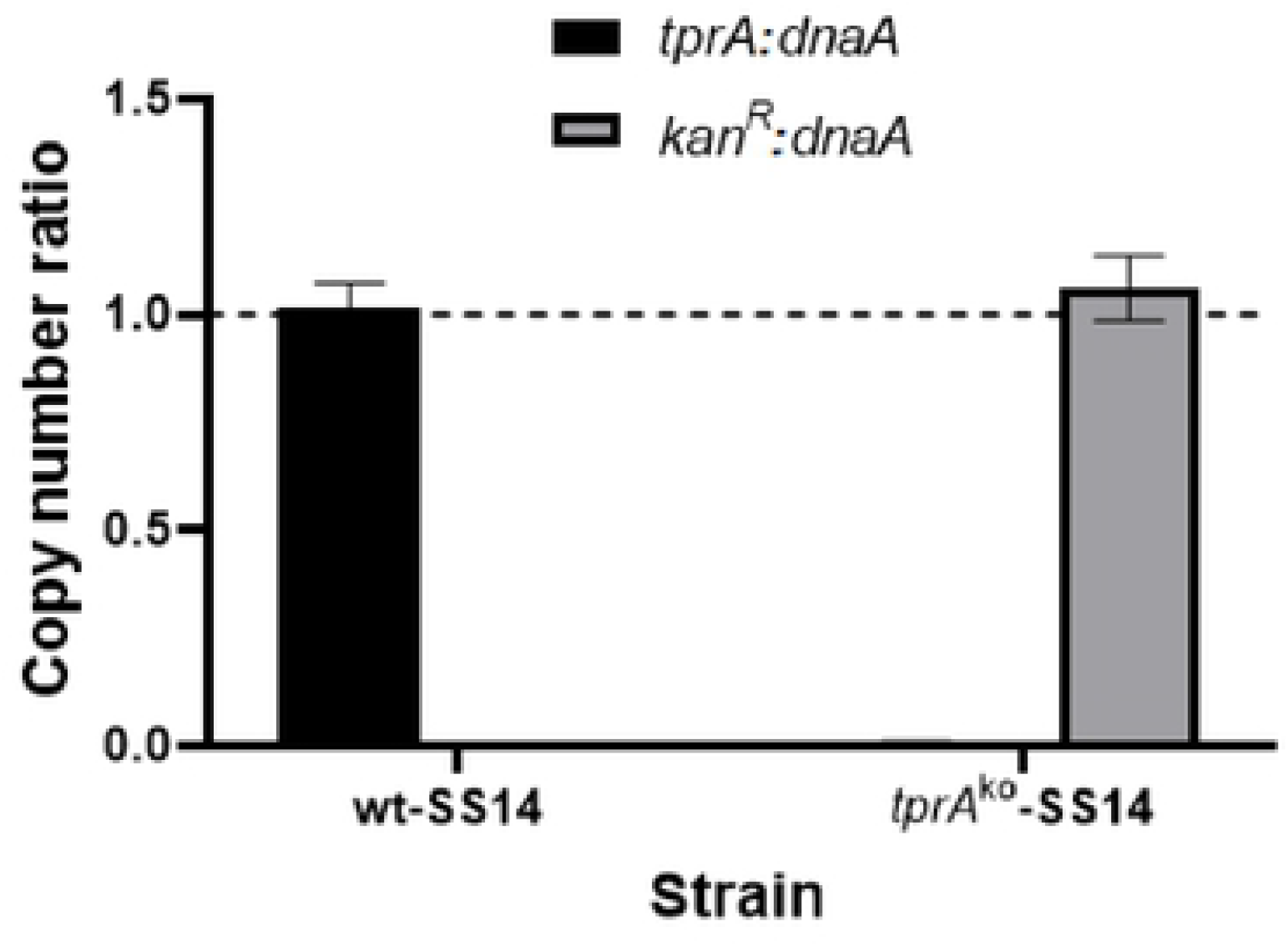
Droplet digital PCR (ddPCR) on DNA template from the *tprA*^ko^- and wt-SS14 strain harvested at Passage #8 post-transformation showing rations between the *kan*^R^, *dnaA*, and *tprA* targets. *tprA*:*dnaA* ratio for the *tprA*^ko^-SS14 strain was 0.006. The *kan*^R^:*dnaA* ratio for the wt-SS14 was zero.

### Whole-genome sequencing from *in vitro*-propagated *tprA*^ko^-SS14 treponemes

At Passage #9 (Week 14 post-transformation; Fig.1), treponemes were harvested for whole-genome sequencing (WGS) using a custom hybridization capture panel to enrich for *T. pallidum*. Genome sequencing showed the absence of the *tprA* ORF in the *tprA*^ko^-SS14 strain when read assembly was performed using the wt-SS14 genome (Fig.4A). However, when assembly used a template genome where *tprA* was replaced with the *kan*^R^ gene, results showed replacement of *tprA* with the *tp0574*promoter-*kan*^R^ sequence (Fig.4B). The wild-type SS14 propagated in parallel to the knockout strain was also sequenced as a control. In the wt-SS14, the *tprA* locus is intact (Fig.4C), while a gap appeared when reads from the wt-SS14 strain were assembled to the *tprA*^ko^-SS14 strain genome with *kan*^R^ in place of *tprA*. Because our sequencing approach used enrichment probes based on multiple wild-type *T. pallidum* genomes and did not include probes for the *kan*^R^ gene, coverage of the *kan*^R^ gene (Fig.4B) was slightly lower than the average for the other regions of the *T. pallidum* genome (Fig.4B). These results showed again that the *kan*^R^ gene replaced the *tprA* locus in the *tprA*^ko^-SS14 strain. A search for reads matching to the plasmid backbone was also conducted but yielded no mapping reads, showing lack of residual plasmid in the culture.

**Figure 4.**
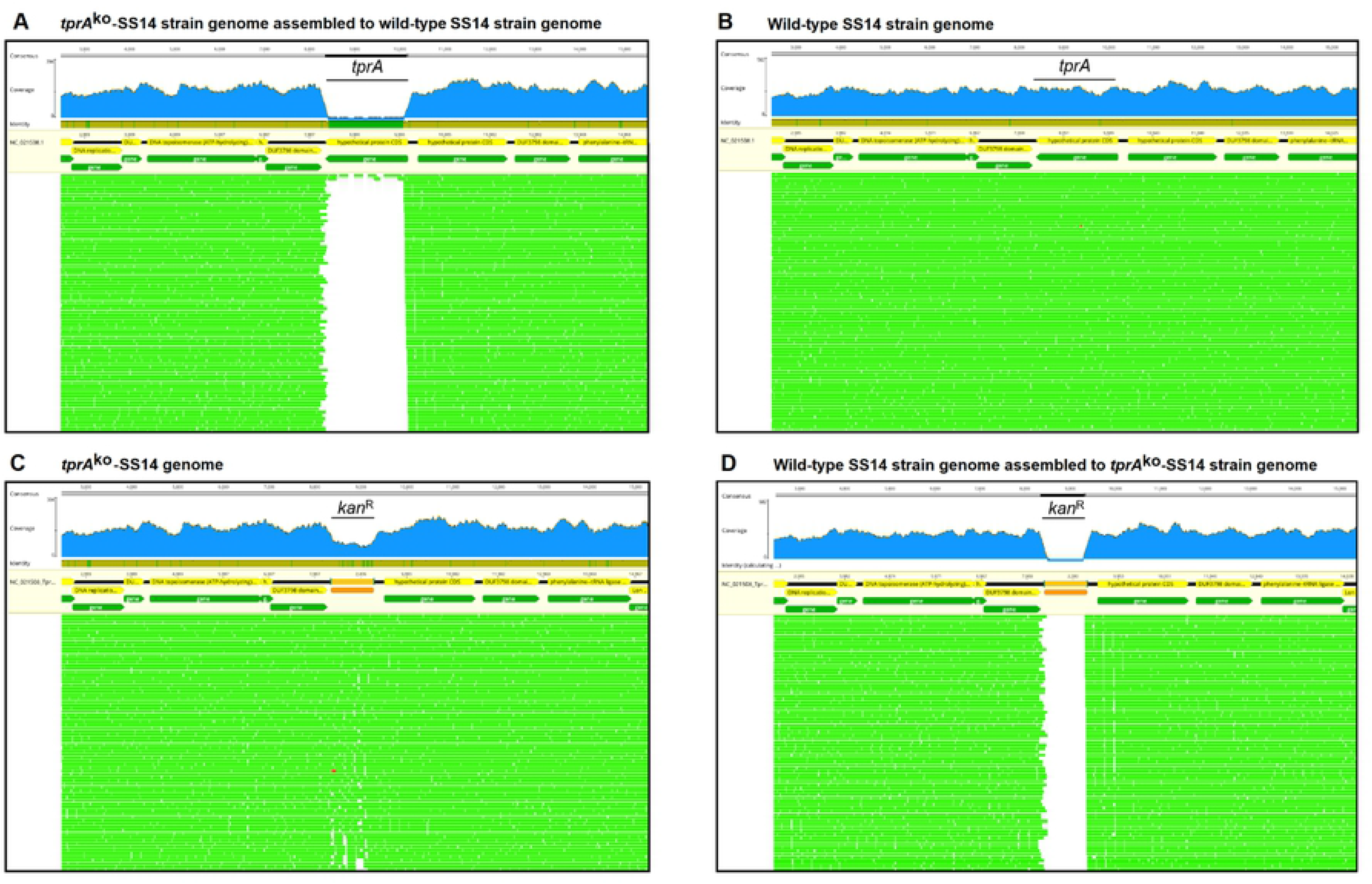
Whole-genome sequencing of DNA template from the *tprA*^ko^- and wt-SS14 strain harvested at Passage #9 post-transformation. (**A**) *tprA*^ko^-SS14 reads assembled to the wt-SS14 genome sequence (NC_021508.1/CP004011.1) showing a gap where the *tprA* locus previously was. (**B**) *tprA*^ko^-SS14 reads assembled to the wt-SS14 genome sequence where the *tprA* locus was replaced *in silico* with the *tp0574*promoter-*kan*^R^ sequence showing reads aligning to the *tp0574*promoter-*kan*^R^ sequence. (**C**) Reads from the wt-SS14 genome (sequenced here as control) aligned to the SS14 reference genome (NC_021508.1/CP004011.1) showing the integrity of the *tprA* locus. (**D**) Reads from the wt-SS14 genome (sequenced here as control) aligned to the *tprA*^ko^-SS14 genome showing a gap where the *tp0574*promoter-*kan*^R^ sequence is located.

### Kanamycin susceptibility assay

*tprA*^ko^-SS14 were grown in 25 µg/ml of kanamycin during routine propagation, a concentration shown to be treponemicidal for the wild-type (Fig.1). To demonstrate that the *tprA*^ko^-SS14 strain could grow at significantly higher kanamycin concentration, *tprA*^ko^-SS14 cells harvested at Passage #10 (Week 15 post-transformation; Fig.1) were used to further assess resistance to kanamycin by performing an *in vitro* susceptibility assay. To this end, we grew both the wt-SS14 and *tprA*^ko^-SS14 strains in media supplemented with kanamycin ranging from 200 to 1.6 μg/ml, for a total of 8 different concentrations tested in 8 replicate wells. Quantification of treponemal burden, measured by qPCR targeting the *tp0574* gene showed that media supplemented with 200 down to a MIC of 3.13 µg/ml of kanamycin strongly inhibited the growth of the wild-type strain when compared to no-antibiotic wells (Fig.5A), but did not affect the *tprA*^ko^-SS14 strain, which grew as if no antibiotic was added (Fig.5B), thus confirming that *tprA*^ko^-SS14 treponemes had become resistant to kanamycin. Furthermore, the *tprA*^ko^-SS14 strain also appeared to have a growth advantage compared to the wild-type strain. When growth was compared at day 7 post-inoculation using the no-antibiotic wells, the *tprA*^ko^-SS14 was shown to have grown significantly faster than the wild-type, with an average of ∼12,000 genome copies/µl, compared to the ∼4,300 copies of the wild-type strain (*p*<0.05), even though the initial inoculum size was the same. As a control, DNA extracted from the eight *tprA*^ko^-SS14 replicate cultures grown in 200 µg/ml of kanamycin was amplified using primers specific for the backbone of the pUC57 plasmid (pair #5/6, as shown in Fig.2A), to ensure no residual plasmid was present in these cultures. All these amplifications yielded a negative result unless the p*tprA*arms-*tp0574*prom-*kan*^R^ construct was used as positive control (Fig.5C), confirming no presence of residual plasmid in these cultures. When the same DNA was tested with primers flanking the *tprA* homology arms, the *tprA*^ko^-SS14 cultures yielded a ∼3.7Kbp amplicon, expected due to the replacement of the *tprA* locus with the shorter *kan*^R^ sequence, while the wt-SS14 strain DNA yielded a ∼4.6 Kbp amplicon (Fig.5D).

**Figure 5.**
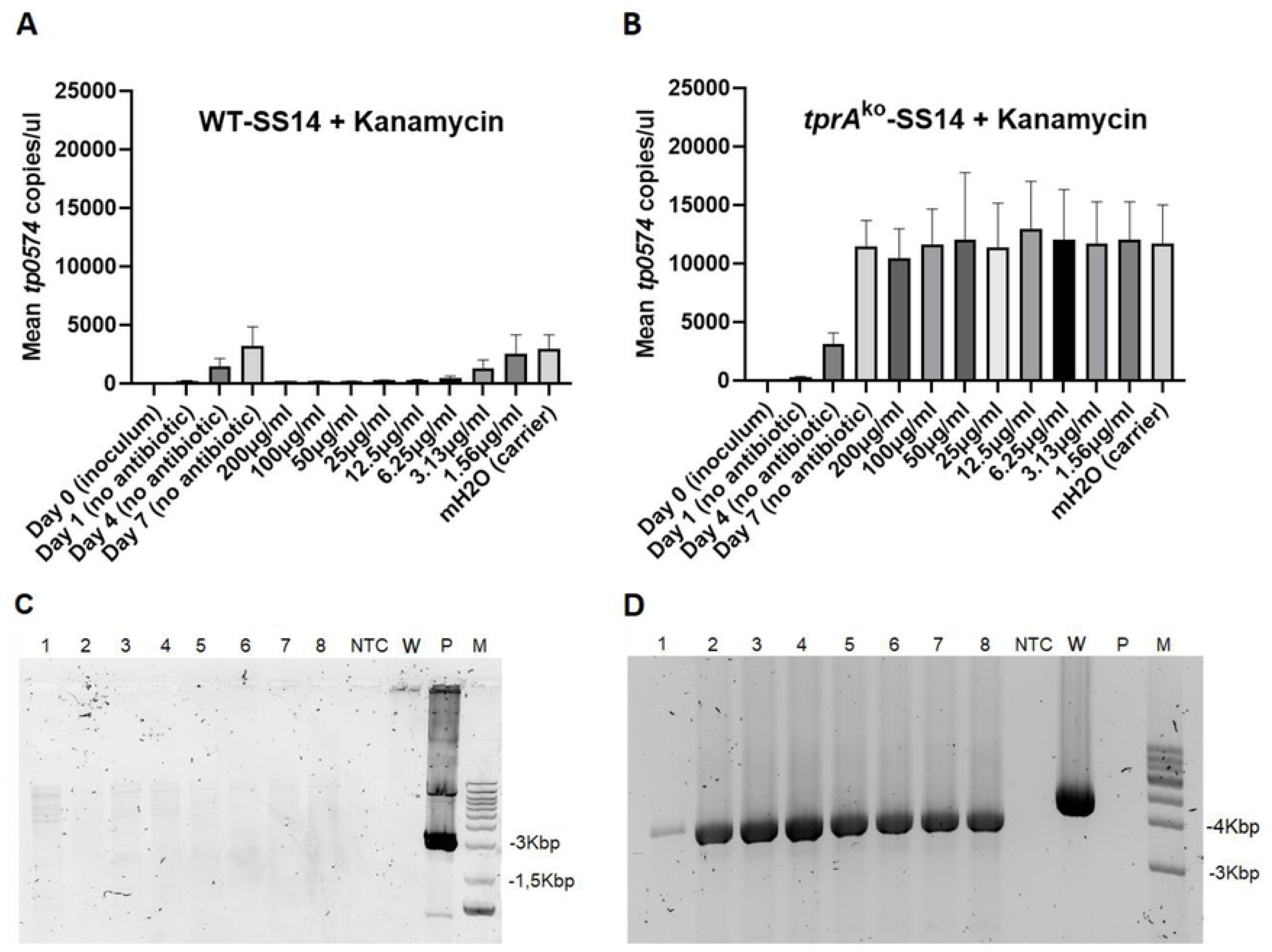
Kanamycin susceptibility assay for the wt-SS14 strain (**A**) and the *tprA*^ko^-SS14 (**B**). In panel A, values (mean ± SD) for Day 0 and Day 1 are 8.95 (±1.6) and 209.6 (±12.2), respectively. In panel B, values for Day 0 and Day 1 (mean ± SD) are 18.0 (±1.8) and 315.6 (±14.7), respectively. mH2O: molecular grade water. DNA was extracted from the 8 replicate *tprA*^ko^-SS14 cultures propagated in 200 µg/ml of kanamycin and tested with primers targeting the pUC57 vector backbone (panel **C**) used for transformation, showing lack of amplification. Wells 1-8: DNA template from the 8 replicate cultures; NTC: no template control; N: wt-SS14 strain DNA; P: p*tprA*arms-*tp0574*prom-*kan*^R^ plasmid DNA template; M: molecular size marker. Expected amplicon size in ∼3Kb. (**D**) DNA was extracted from the 8 replicate *tprA*^ko^-SS14 cultures propagated in 200 µg/ml of kanamycin and tested with primers targeting *T. pallidum* genomic region right outside of the *tprA* homology arms (primers in Table 1), showing that in the *tprA*^ko^-SS14 strain a smaller amplicon is obtained due to the replacement of *tprA* by the *kan*^R^ gene, which is approximately 1Kb shorter in size. Wild-type SS14 DNA template yielded a longer amplicon due to an intact *tprA* locus still in place. No amplification was detected using the transformation plasmid DNA control. NTC: no template control; W: wt-SS14 strain DNA; P: p*tprA*arms-*tp0574*prom-*kan*^R^ plasmid DNA template; M: molecular size marker (Kbp).

The ratio *kan*^R^:*dnaA* in the *tprA*^ko^-SS14 cultures grown at different kanamycin concentrations estimated by ddPCR ranged between 1.07 and 1.14 (Fig.6A) on average. On the contrary, in wt-SS14 the *kan*^R^:*dnaA* ratio was zero. In the *tprA*^ko^-SS14 cultures, the *tprA*:*dnaA* ratio was virtually zero (0.004; Fig.6B), while the *tprA*:*dnaA* ratio was 1.18 on average (Fig.6B). The ratio *kan*^R^:*tp0574* in *tprA*^ko^-SS14 grown in different kanamycin concentrations, was shown to be in average 1.40 in treponemes analyzed after 7 days in culture, and slightly higher (1.66) in treponemes harvested after 4 days in culture (Fig.6C), although this difference was not significantly different. This result suggested that more copies of the *kan*^R^ gene than the *tp0574* gene were present in the extracted DNA at sample harvest. This was overall an expected result. The replacement of the *tprA* pseudogene (*tp0009*) in the SS14 strain, in fact, positioned the *kan*^R^ gene in proximity (within 10 Kbp) of *T. pallidum dnaA*, *dnaN* (*tp0002*), and *gyrA* (*tp0005*) genes that, in prokaryotes, are markers for the chromosomal origin of replication (*oriC*) [17, 18]. The >1 ratio *kan*^R^:*tp0574* in *tprA*^ko^-SS14 grown in culture likely reflects partial replication of some chromosomes during propagation, and not differences in amplification efficiency. Such conclusion is also supported by the evidence that in the wild type strain the average *tprA*:*tp0574* ratio, obtained using the same samples above, is 1.31. Overall, these results supported that a) the *kan*^R^ gene is stably integrated into *T. pallidum* genome, that b) there are no residual copies of the *kan*^R^ gene present in episomes, that c) no transformation plasmid is still present and that, for future experiments, d) the copy number of a transgene needs to be compared to that of a neighboring gene to account for replication-induced bias, particularly if the transgene is close to *oriC*.

**Figure 6.**
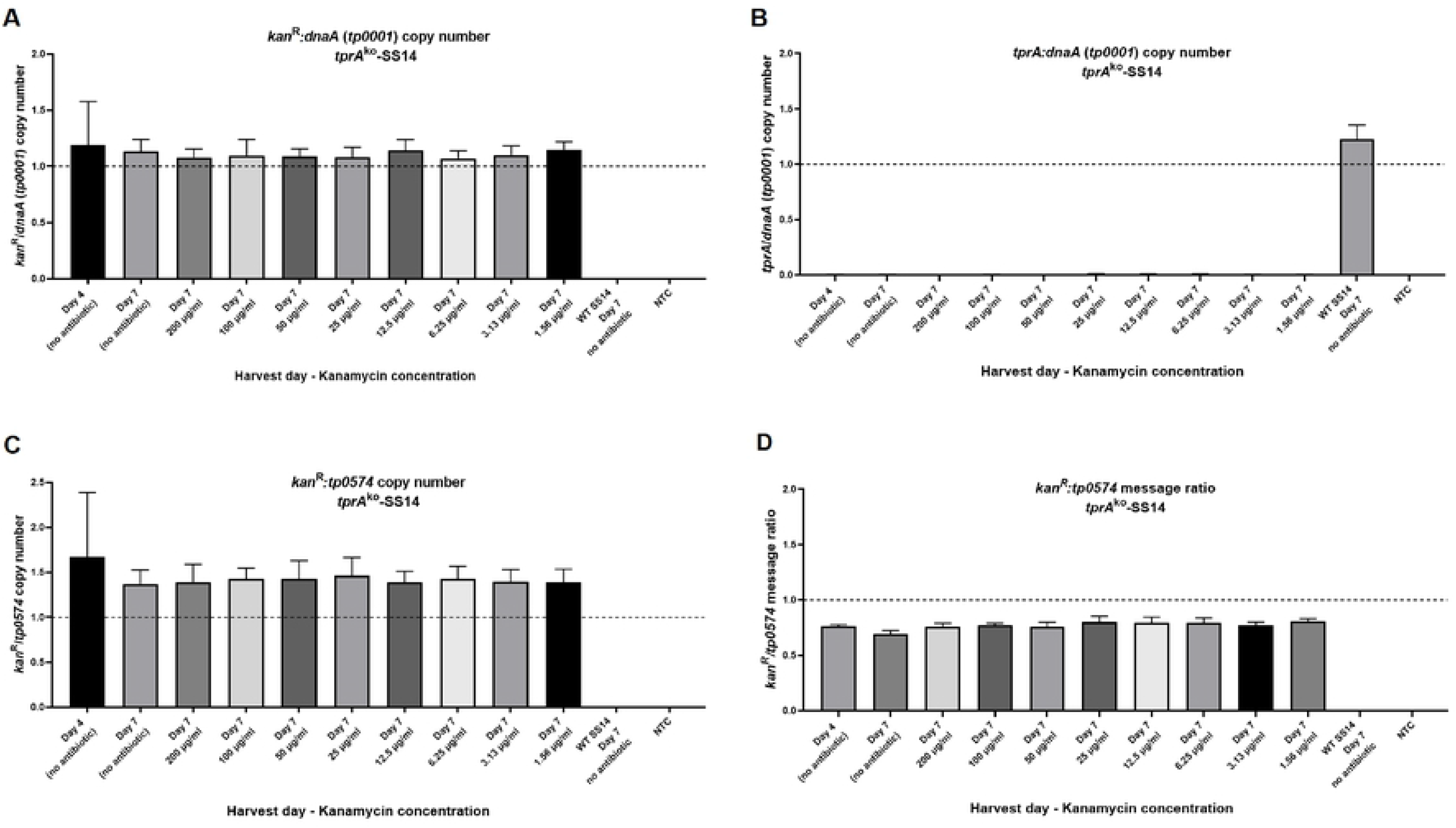
Droplet digital PCR (ddPCR) on DNA or cDNA template from the *tprA*^ko^- and wt-SS14 strain propagated in different kanamycin concentrations (*tprA*^ko^-SS14) or no antibiotic (wt-SS14). (**A**) *kan*^R^:*dnaA* gene copy number ratio. (**B**) *tprA*:*dnaA* gene copy number ratio. (**C**) *kan*^R^/*tp0574* gene copy number ratio. (**D**) *kan*^R^:*tp0574* mRNA ratio. NTC: no template control.

Because in the knockout strain the *kan*^R^ gene and the *tp0574* gene are transcribed by the same promoter, cDNA was used to quantify the message level for these two genes. The *kan*^R^:*tp0574* message ratio was found to be 0.77, which showed the *tp0547* gene being slightly more highly expressed than the *kan*^R^ gene. This result suggested that the choice of using the *tp0574* promoter to drive expression of the *kan*^R^ gene led to very similar message levels for these genes, as hypothesized during the experimental design of the p*tprA*arms-*tp0574*prom-*kan*^R^ plasmid (Fig.6D). Amplification of *kan*^R^ message was not detected in the wt-SS14 strain, and no *tprA* message amplification occurred when cDNA from the knockout strain was used as the template. *tprA* message was however detected by ddPCR in the wt-SS14 strain. In this case the *tprA*:*tp0547* ratio was 0.040, confirming that the level of transcription of the *tprA* pseudogene is extremely low, compared to *tp0547*.

### Rabbit infection

We next examined how the *tprA*^ko^-SS14 strain acted during *in vivo* infection in the New Zealand white rabbit model, expecting that the strain would survive kanamycin treatment of the animal. The rabbit infected with the *tprA*^ko^-SS14 strain developed orchitis of the left testicle on day 17 post-inoculation. Treponemal yield from the animal was 1.2×10^8^ *T. pallidum* cells/ml of testicular extract. At the time of harvest, the animal was seropositive with the *Treponema pallidum* particle agglutination test (TPPA) and the Venereal Disease Research Laboratory (VDRL) test, confirming the establishment of infection. The control rabbit, infected with the wt-SS14 strain but not treated, developed orchitis at day 24 post-infection, and the treponemal yield was 2.2×10^7^ *T. pallidum* cells/ml of testicular extract. This animal was also TPPA-positive but VDRL-negative. On day 24 post-infection, the control rabbit infected with the wt-SS14 strain and subcutaneously treated with kanamycin had not developed orchitis and was euthanized (repeat intramuscular injection of kanamycin was not allowed by local IACUC). Upon analysis of the testicular exudate from this animal, treponemes could be seen, suggesting that the subcutaneous treatment with kanamycin was not completely effective *in vivo* as it was *in vitro* (Fig.1). From this animal, the treponemal yield was however much lower compared to the other rabbits (1.5×10^5^ *T. pallidum* cells/ml, based on detecting 3 treponemal cells in 20 DFM fields). As a further confirmation of treatment failure, this animal was also TPPA-positive but VDRL negative. These data suggested that, although the kanamycin concentration achieved in the rabbits via subcutaneous injection was not completely treponemicidal, *tprA*^ko^-SS14 were less susceptible to the antibiotic and proliferated faster than the wild-type. DNA extracted from these treponemal harvests was used for ddPCR targeting the *dnaA*, *tprA*, and *kan*^R^ genes. Droplet Digital PCR showed that for the *tprA*^ko^-SS14 propagated *in vivo*, the *kan*^R^:*dnaA* and *tprA*:*dnaA* ratios were 1.05 and 0.007, respectively (Fig.7). For the wt-SS14 strain extracted from the untreated and treated rabbits, respectively, the *kan*^R^:*dnaA* and *tprA*:*dnaA* ratios were 0.00 and 1.02, respectively (untreated rabbit) and 0.00 and 0.83, respectively (ineffectively treated rabbit; Fig.7). In these samples the *kan*^R^:*tp0574* ratio was also obtained and was found to be 1.30 (*tprA*^ko^-SS14), and 0.000 (wt-SS14, in both the untreated and ineffectively treated animals. All these *in vivo* ddPCR results are consistent with ratios seen during *in vitro* propagation (Fig.3 and Fig.6).

**Figure 7.**
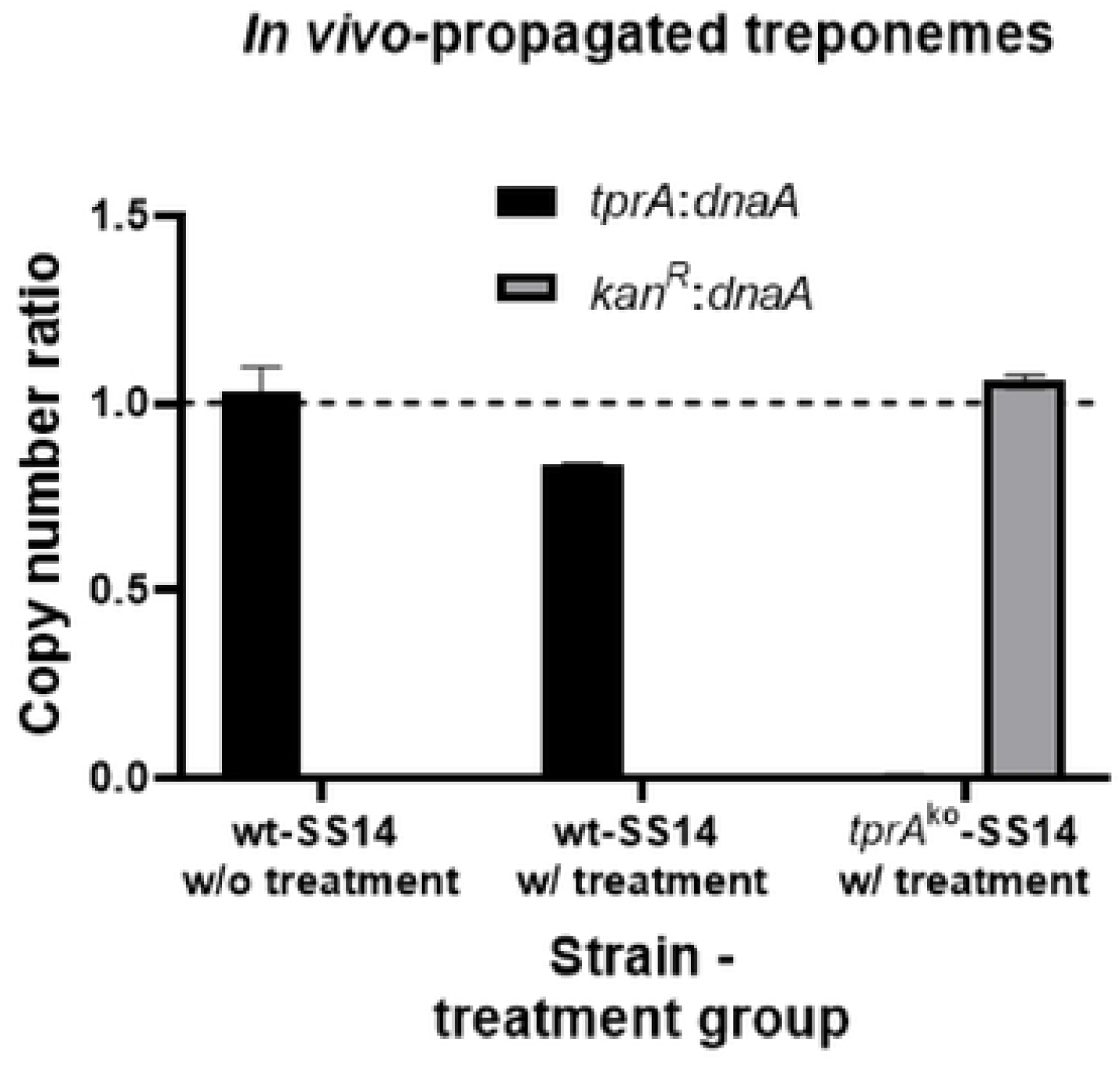
Droplet digital PCR (ddPCR) on DNA template from the *tprA*^ko^- and wt-SS14 strain propagated in rabbits to which kanamycin was (w/) or was not (w/o) given. The only IACUC-approved administration route for kanamycin, however, resulted in treatment failure in rabbits infected with the wt-SS14 strain. *tprA*:*dnaA* ratio for the *tprA*^ko^-SS14 strain was 0.007; *kan*^R^:*dnaA* ratio for the wt-SS14 strains was zero.

### Mass spectrometry

To further confirm the expression of the 31.01 kDa Kan^R^ protein, we performed liquid MS on proteins of the *tprA*^ko^-SS14 and wt-SS14 strains separated by SDS-PAGE and ranging approximately from 20-45 kDa. This portion of the *T. pallidum* proteome was retrieved through of band excision from the acrylamide gel to then undergo in-gel tryptic digestion before MS, and the size range (20-45 kDa) was decided since a distinct ∼31 kDa band corresponding to Kan^R^ could not be undoubtedly identified in the Coomassie-stained gel. Nonetheless, MS data analysis showed that peptides mapping to 77% of the Kan^R^ protein could be isolated from the *tprA*^ko^-SS14 strain sample (Fig.8), but not from the paired sample from the wild-type strain. Based on the MS, the amount of the Kan^R^ protein corresponded to ∼1% of the total protein content detected in the specimen. The list of all peptides mapping to the Kan^R^ protein is reported in Table 2. The full MS results for the samples corresponding to the *tprA*^ko^-SS14 and wt-SS14 are provided as Supporting Information (files labeled SS14_TprA_KO, and SS14_WT, respectively). In the SS14_TprA_KO file, Kan^R^ peptides are labeled PMC_FU_2039. These data demonstrated the expression of the Kan^R^ protein in the *tprA*^ko^-SS14 following integration of the gene. No peptides corresponding to the TprA protein (translated using both the +1/+2 reading frames to overcome the effect of the frameshift) were found in the *tprA*^ko^- or wt-SS14 samples.

**Figure 8.**
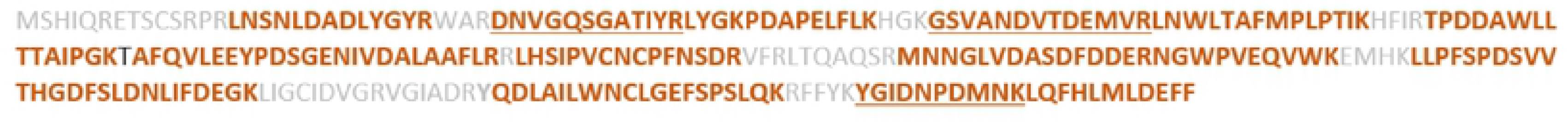
Mass spectrometry coverage of the Kan^R^ protein expressed by the *tprA*^ko^-SS14 strain. Orange sequences were experimentally identified. Light gray sequences were not. Protein coverage is 77%. If two identified peptides are adjacent, one is underlined. Overall, the Kan^R^ protein represented 1% of all proteins identified in the specimen.

**Table 2.**
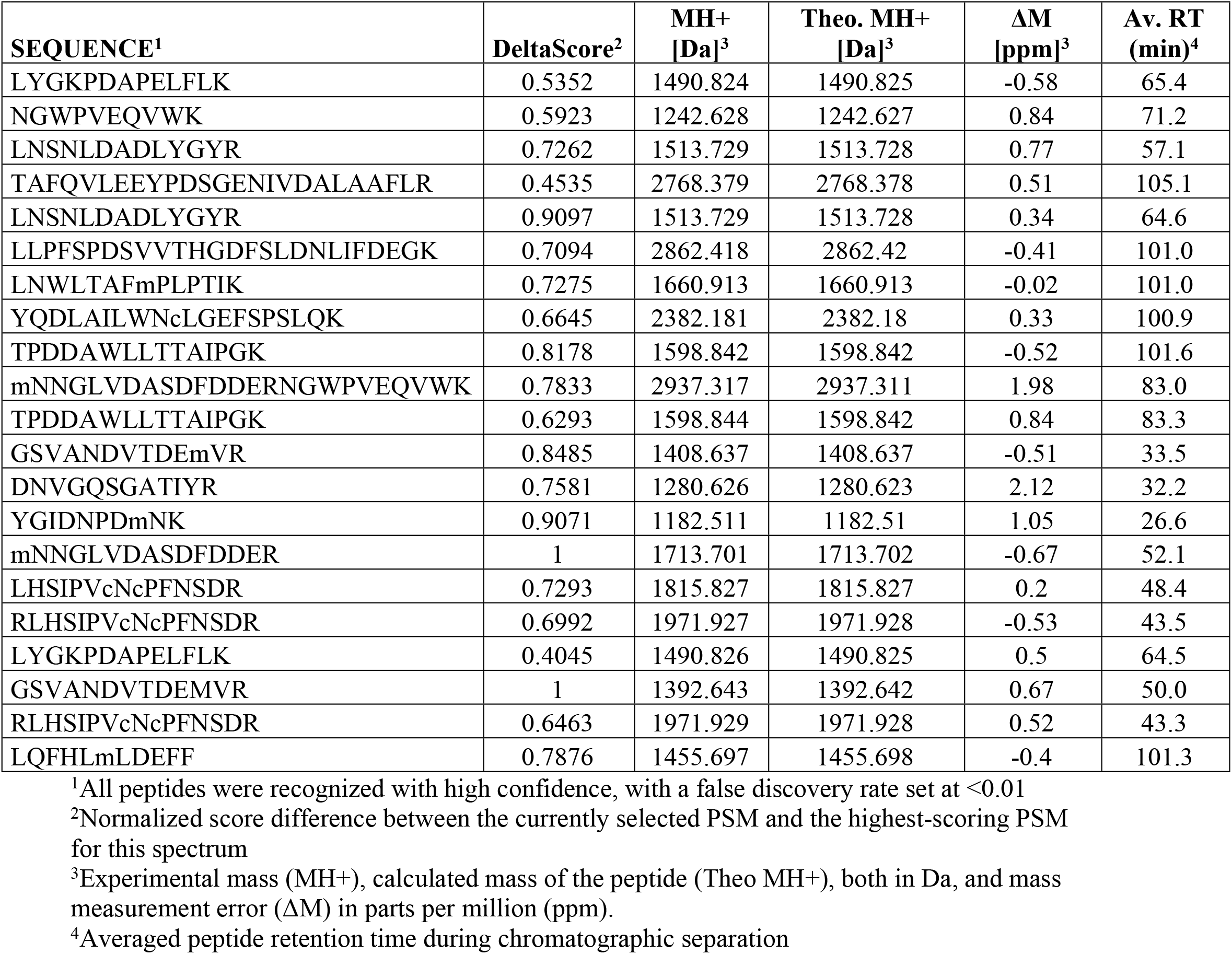
Peptides identified by MS on the tprA^ko^-SS14 strain matching the Kan^R^ protein sequence.

### Derivation of a *tprA*^ko^-SS14 pure isolate by limiting dilution and increased antibiotic pressure

The *tprA*^ko^-SS14 inactivates kanamycin by expressing an aminoglycoside N6’-acetyltransferase that that catalyzes the conversion of kanamycin to N6’-acetylkanamycin using acetyl-CoA. We reasoned that if enough *tprA*^ko^-SS14 cells were present, they could bring kanamycin concentration below the minimal inhibitory concentration for this antibiotic and allow growth of wild-type cells. To eliminate the possibility that wt-SS14 *T. pallidum* cells might survive among the *tprA*^ko^-SS14 cells, we performed serial dilutions of the *tprA*^ko^-SS14 cells and plated them on multiple wells of a 96-well plate containing Sf1Ep cells. Specifically, wells were seeded with 3300 and 330 treponemal cells from the *tprA*^ko^-SS14 culture. Additionally, we increased kanamycin concentration to 200 µg/ml, shown already to be ineffective against the *tprA*^ko^-SS14 strain (Fig.5A). Although these cultures were passaged every two weeks to allow treponemal growth, exhausted media was exchanged once a week to ensure Sf1Ep cells viability and replenish the kanamycin supply. Lack of wt-SS14 cells was assessed by performing ddPCR targeting the *tprA*, *kan*^R^, and *dnaA* genes as described in the Methods section. Results (Fig.9) showed that, after two (Fig.9; P1) and four (Fig.9; P2) weeks in culture, no *tprA*-specific signal could be obtained by ddPCR, supporting complete lack of wild-type strain in the culture wells, and that in the *tprA*^ko^-SS14 strain, the *kan*^R^:*dnaA* copy number was virtually identical.

**Figure 9.**
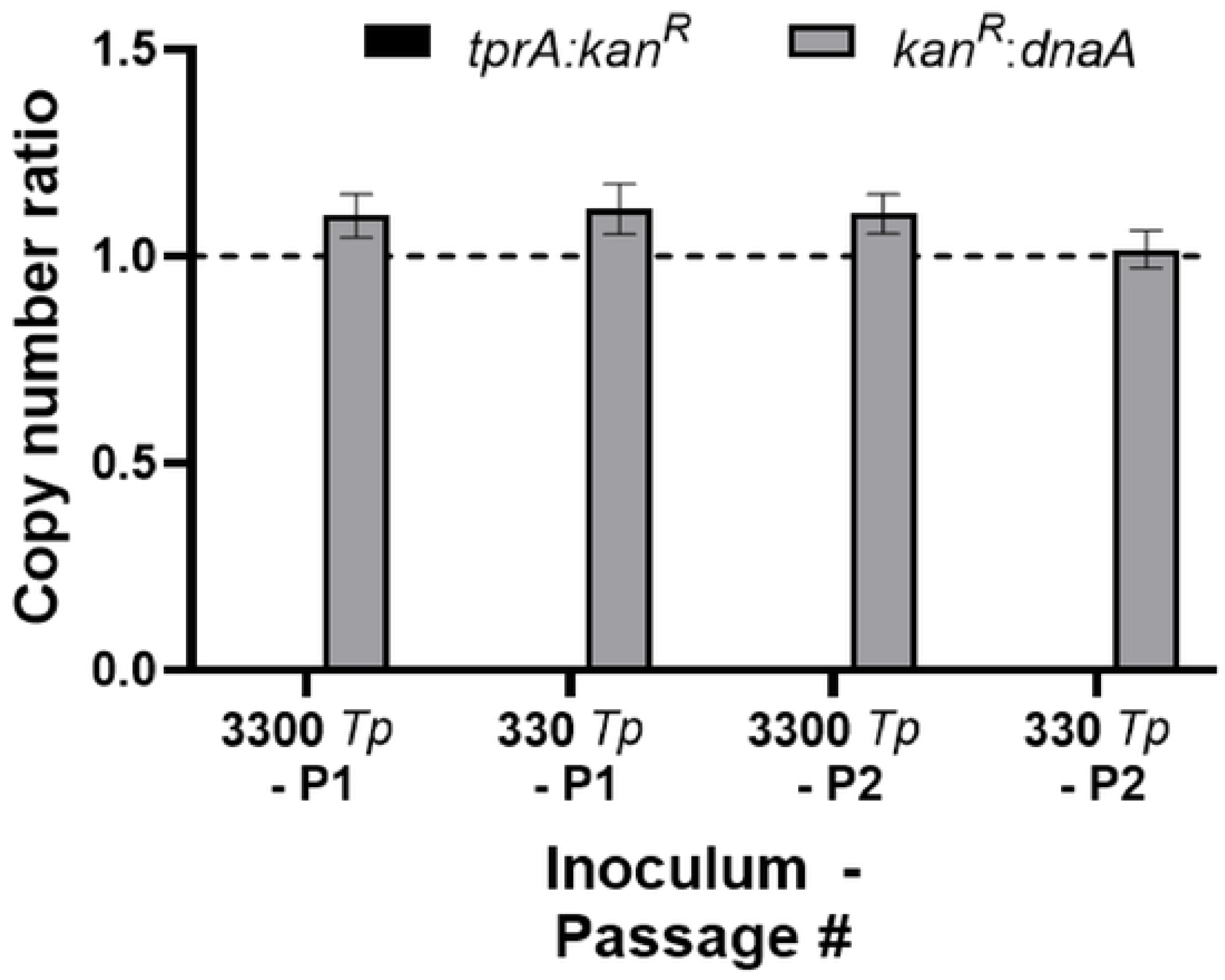
Expansion of the *tprA*^ko^-SS14 culture after inoculation of a limited number of treponemal cells. Two weeks past inoculation (P1), no *tprA*-specific signal could be detected by ddPCR, while the ratio *kan*^R^:*dnaA* was virtually 1. The same results were obtained four weeks after inoculation (P2).

## Discussion

Given the significant impact of syphilis on human health, there is great interest in deepening our understanding of the molecular mechanisms underlying its pathogenesis. In turn, this could lead to improved strategies for disease control and vaccine development. A critical step in this direction is the identification of treponemal factors that contribute to virulence. To date, several experimental approaches have helped syphilis investigators identify and functionally characterize these factors, such as expression in heterologous hosts (e.g. *Borrelia burgdorferi*, *Treponema denticola*, and *Treponema phagedenis*) [19–21], comparative genomics [22–26] and, to some extent, gene expression studies and proteomic analysis [15, 25, 27]. Genetic manipulation strategies, however, particularly those that satisfy Koch’s molecular postulates [28], have not been available for *T. pallidum*. The lack of genetic tools was not overly frustrating because, until 2018 [11, 27], *T. pallidum* could not even be continually propagated *in vitro*.

Understandably, this limitation hindered any attempt aimed at genetically engineering this pathogen, as antibiotic selection could not readily be performed. In contrast, some *Chlamydia* species have been propagated *in vitro* since the 1950s, and yet genetic tools for this pathogen became available only recently [11, 29]. The protocol we devised to make our vector cross the *T. pallidum* envelope was indeed inspired by an early protocol used to introduce DNA into *Chlamydia* [22, 29]. The use of CaCl_2_ was preferred to that of physical methods such as electroporation simply because the number of treponemal cells yielded by *in vitro* culture is still very limited compared to other bacteria, including other spirochetes, that have a generation time significantly shorter than *T. pallidum* and that can be propagated *in vitro* in axenic cultures. Years ago, before the introduction of the *in vitro* cultivation system for *T. pallidum*, our laboratory considered an alternative approach to introducing foreign DNA into *T. pallidum.* Despite *T. pallidum’s* lack of any known plasmids or phage, the mounting evidence that inter-strain recombination occurs in *T. pallidum* strains and subspecies [22, 23, 30], prompted us to hypothesize that *T. pallidum* could be competent for transformation. Back then, to test this hypothesis, we exposed Nichols cells, shortly after harvest from rabbit testes to a construct similar to the one used here, but with a chloramphenicol resistance (instead of *kan*^R^) gene under control of the *tp0574* promoter, between *tprA* homology arms. The construct also contained several DNA uptake sequences (DUSs) [31] from both Gram-positive and Gram-negative bacteria to facilitate uptake by a putative surface receptor/internalization machinery that was predicted *in silico* in *T. pallidum* based on comparative genomics. Following exposure to the construct, treponemes were re-injected into a naïve animal’s testes, and selection was attempted *in vivo* by treating the animal with chloramphenicol. Although these previous experiments failed to provide evidence that genetically modified treponemes could be obtained, as treponemes could be not retrieved from the infected animal despite seroconversion, the hypothesis that *T. pallidum* could naturally uptake DNA from the environment and integrate it into its genome can now be explored again, to help explain inter-strain recombination events.

New and far more exciting future directions include the development of shuttle plasmids that do not need to integrate into the *T. pallidum* genome but are suitable to express *T. pallidum* ORFs for complementation purposes, for example, but also to express fluorescent reporter proteins such as GFP, CFP, mCherry, or reporter enzymes such as β-galactosidase, with the overall goal of better understanding gene regulation. Expression of luciferase enzymes for *in vivo* imaging, as done for other spirochetes [20, 30], could also be desirable. The transformation of *T. pallidum* strains with shuttle plasmids should not be complicated by the presence of native plasmids like it is in some *Chlamydia* species. Plasmids carrying the same origin of replication are generally incompatible and tend not to coexist in a cell. Therefore, the maintenance of a native plasmid limits the introduction of exogenous ones [20, 32]. The absence of endogenous plasmids in *T. pallidum* should eliminate the issue of competition for plasmid replication factors that could hinder the transformation efficiencies of exogenous recombinant plasmids.

Also, the development of a conditional expression vector for proteins or miRNA for RNA silencing and post-transcriptional regulation would be highly beneficial. Our first attempt to knock out the *tprK* gene (*tp0897*) of *T. pallidum* was conducted in parallel to the experiments described in this manuscript. We were, however, unsuccessful in obtaining a *tprk*^ko^ strain. TprK is a *T. pallidum* outer membrane protein (OMP) that undergoes extensive intra-strain antigenic variation through non-reciprocal gene conversion and is one of the virulence factors primarily responsible for *T. pallidum* immune evasion during infection [32–35]. Despite the extensive recombination events that affect the gene ORF, *tprK* gene variants with frameshift mutations or stop codons were never reported, even when deep sequencing was performed on this gene. This could suggest that *T. pallidum* cells with a non-functional TprK are not viable and is consistent with the spirochete’s need to generate extreme diversity in this gene rather than simply deleting it. Regarding our experiment to replace *tprK* with a *kan*^R^ cassette, we might simply have been unsuccessful. However, if a *tprK*^ko^ strain were not viable, a conditional expression system could circumvent the problem and be useful to understand what biological function the TprK protein variants mediate in addition to immune evasion that makes it essential for this pathogen. In the specific case of TprK, however, current work is aiming at deleting the source of *tprK* variability, namely the donor cassettes that recombine into the gene expression site. This should generate treponemes with an impaired recombination system that can only express a single TprK variant, hence incapable of immune escape. Along the same line, the inactivation of many genes might be problematic in the syphilis spirochete, because the small size of treponemal genomes suggests that the genes that escaped evolutionary genomic reduction might be essential. In this scenario, inducible systems for gene silencing or ablation would be helpful. Today, site-directed recombination technologies are increasingly used to manipulate an organism’s DNA under controlled conditions *in vivo*. One example is the system analogous to Cre-Lox recombination but involves the recombination of sequences between short flippase recognition target (FRT) sites by the recombinase flippase (Flp). An FRT-Flp recombination system would allow *in vivo* gene ablation if FRT could be introduced upstream and downstream of a locus. Expression in *trans* of the Flp recombinase would then induce excision of the sequence between the FRTs. In addition, an inducible expression system would be beneficial for epigenetic gene silencing approaches, such as the CRISPR/Cas9 system and TALENS [35–38]. All these approaches, however, will require some adjustments to the biology of *T. pallidum*, whose genome has a very high GC content (∼56%), and certain genes expressing ectopic proteins such as recombinases will need to be optimized for expression in this spirochete. A different direction could take advantage of transgenic expression to increase the amount of OMPs in *T. pallidum* envelope in a *tprK*-impaired mutant. This approach could lead to a strain that could be used to create outer-membrane vesicles rich in protective OMPs to be used for immunization purposes.

The result that the ratio between the *kan*^R^ gene and *tp0574* was higher than originally expected was intriguing and worth discussing in the context perhaps of molecular detection assays for *T. pallidum*. Following the sequencing of the Nichols strain genome, *T. pallidum oriC* was localized at nucleotide +1 [39] based on gene synteny. Replication of the bacterial chromosome is initiated at a single *oriC* region and proceeds in both directions. During growth, replication is generally initiated once per cell cycle, however, under optimal nutrient and media conditions, another round of replication can be initiated even before the previous round has completed, resulting in the inheritance by daughter cells of partially replicated chromosomes [18, 40, 41]. Partial replication of a chromosome can create an unbalanced ratio between genes that are near *oriC* (such as *kan*^R^, in our case) that are already duplicated, and genes farther away from *oriC* that are still in single copy (such as *tp0574*, located at the polar opposite of the *kan*^R^ gene in the ∼1Mb *T. pallidum* chromosome, and therefore one of the last genes to replicate). Hence, during active cell growth, the ratio between copies of genes close to *oriC* and more distant ones can be remarkably above 1. As mentioned, the >1 ratio *kan*^R^:*tp0574* in *tprA*^ko^-SS14 grown in culture likely reflects partial replication of some chromosomes during propagation. This could support that genes located near the origin of replication could be better targets for *T. pallidum* detection in clinical samples, but perhaps that genes located further away from the origin could be more suitable to estimate the actual number of treponemal cells in a sample.

Ongoing work in the laboratory is focusing also on analyzing the transcriptional and proteome profile of the *tprA*^ko^-SS14 strain compared to the wild-type. It is intriguing that, based on our *in vitro* cultivation results, the *tprA*^ko^-SS14 strain grew significantly faster than the wild-type strain. This difference was not due to errors in the initial inoculum, as the experiment reported in Fig.3 was repeated twice independently and the results were not different. The ability of the transformed strain to proliferate faster was also suggested by the fact that the rabbit infected with the *tprA*^ko^-SS14 strain developed orchitis earlier than the untreated control, even though the inoculum was the same. However, the *in vivo* experiments are not conclusive in this regard, as only one rabbit was used as untreated control, and the later development of orchitis could simply be due to rabbit-to-rabbit variability. It is nonetheless possible that ablating the *tprA* pseudogene might have conferred a selective advantage and have reduced the metabolic burden of expressing a gene that is not functional. Previous studies carried on in our laboratory with *T. pallidum* strains carrying a frame-shifted *tprA* gene (although not with the SS14 strain), showed that *tprA* is transcribed in these strains, although at a very low level [16].

Additional experiments to be performed include repeating the transformation experiment reported here, as well as targeting other *T. pallidum* genes that, unlike *tprA*, will provide us with the ability to study a phenotype. As proteins mediating motility, antigenic variation, and adhesions are main virulence factors of spirochetes, our attention will focus next on ablating expression of the endoflagella and evaluate whether a non-motile *T. pallidum* can successfully establish an infection in the rabbit host, on eliminating the *tprK* donor sites to impair antigenic variation and immune evasion, and knock-out adhesins.

## Conclusions

We demonstrate that genetic engineering of the syphilis spirochete is possible with a relatively simple method that has the potential to “transform” our way to approach the study of *T. pallidum* biology and syphilis pathogenesis.

## Materials and Methods

### Ethics statement

Only male NZW rabbits (*Oryctolagus cuniculus*) ranging from 3.5-4.5 kg in weight were used in this study. Specific pathogen-free (SPF; *Pasteurella multocida*, and *Treponema paraluiscuniculi*) animals were purchased from Western Oregon Rabbit Company (Philomath, OR) and housed at the University of Washington (UW) Animal Research and Care Facility (ARCF). Care was provided in accordance with the procedures described in the Guide for the Care and Use of Laboratory Animals [42] under protocols approved by the UW Institutional Animal Care and Use Committee (IACUC; Protocol # 4243-01, PI: Lorenzo Giacani). Upon arrival and before use, all rabbits were bled and tested with a treponemal test (TPPA; Fujirebio, Tokyo, Japan) and a non-treponemal test (VDRL; Becton Dickinson, Franklin Lakes, NJ) to confirm lack of immunity due to infection with *Treponema paraluiscuniculi*, given that animals are tested randomly by the provider. Both tests were performed according to the manufacturer’s instructions. Only seronegative rabbits were used for experimental infection with transformed and wild-type treponemes (see paragraph below for rabbit infection, treatment and sample collection).

### Plasmid construct

The pUC57 vector (2,710 bp; Genscript, Piscataway, NJ) was engineered to carry the *kan*^R^ gene downstream of the *T. pallidum tp0574* gene promoter and ribosomal binding site. Appropriate spacing (8 nt) was ensured between the RBS and the *kan*^R^ gene start codon in the construct. Upstream and downstream of the *tp0574*prom-*kan*^R^ hybrid sequence, respectively, two homology arms corresponding to the regions flanking the *tprA* gene were inserted. The upstream arm was 998 bp in length and corresponded to position 7,343-8,340 of the wt-SS14 strain genome (NC021508.1/ CP004011.1). The downstream arm was 999 bp, and encompassed position 10,165-11,163 of the SS14 genome. The construct was cloned between the XheI and BamHI sites of the pUC57 vector, in opposite orientation compared to the *lac* promoter that is upstream of the polylinker. Prior to use, the insert underwent Sanger sequencing to ensure sequence accuracy. The sequence of the insert is provided in File S1. This construct was named p*tprA*arms-*tp0574*prom-*kan*^R^. Primers annealing a) to the vector only, b) within the cloned insert, and c) upstream of the *tprA* homology arms in the *T. pallidum* genome are reported in Table 1. The pUC57 vector carries an ampicillin resistance gene (*bla*) for selection in *E. coli*. Because penicillin is the first-line antibiotic to cure syphilis, we first evaluated whether the sequences flanking the *bla* gene of the pUC57 vector could have had sufficient homology to *T. pallidum* DNA to induce recombination and integration of the *bla* gene in the genome, but no homology was found upon BLAST analysis of these regions against the SS14 or other syphilis strain genomes. Regarding the insertion of a *kan*^R^ gene in the *T. pallidum* genome, the CDC STI treatment guidelines do not recommend the use of kanamycin for syphilis therapy, hence a kanamycin-resistant syphilis strain does not pose a risk in case of unlikely exposure. To obtain a highly concentrated, endotoxin-free plasmid preparation, the p*tprA*arms-*tp0574*prom-*kan*^R^ was transformed into TOP10 *E. coli* cells (Thermo Fisher, Waltham, MA), which were then grown first in a 5-ml starter culture overnight, and then in 500 ml of LB media supplemented with 100 µg/ml of ampicillin at 37⁰C. The plasmid was purified using the Endo-Free Plasmid Mega Kit (Qiagen, Germantown, MD) according to the manufacturer’s instructions. Following purification, plasmid concentration was assessed using an ND-1000 spectrophotometer (Nanodrop Technologies, Wilmington, NC). The vector was then divided into 50 µl aliquots and stored at -80 until use.

### Source of *T. pallidum* for *in vitro* cultivation, transformation, and selection

The SS14 strain of *T. pallidum* used for *in vitro* propagation was obtained from a frozen stock previously propagated IT in NZW rabbits as already reported [43]. This strain was originally isolated in 1977 in Atlanta (USA) from a penicillin-allergic patient with secondary syphilis who did not respond to therapy with macrolides. *In vitro* culturing was performed according to Edmondson *et al.* [11] in the wells of a 24-well plate initially, and then expanded in a 6-well culture plate (Corning Inc, Corning, NY). The microaerophilic atmosphere (MA; 1.5% O_2_, 3.5% CO_2,_ and 95% N_2_) necessary to sustain treponemal viability was achieved using a Heracell VIOS 160i tri-gas incubator (Thermo Fisher). Before the addition of the treponemal cells, Sf1Ep cells were incubated in a 5% CO_2_ atmosphere in a HeraCell 150 incubator (Thermo Fisher). For transformation, treponemes were first sub-cultured into the wells of a 24-well plate as per protocol [11]. Briefly, the day before treponemal inoculation, a 24-well plate was seeded with 2×10^4^ rabbit Sf1Ep cells/well in 2.5 ml of culture media. The plates were then incubated overnight in the HeraCell incubator. On the same day, TpCM-2 media was prepared according to protocol and equilibrated overnight at 34°C in the MA incubator. The following day, cell culture media was removed from the 24-well plate, and cells were rinsed with equilibrated TpCM-2 media. Subsequently, each well was filled with 2.5 ml of equilibrated TpCM-2 media, and the plate was transferred to the MA incubator. To prepare the treponemal inoculum, the Sf1Ep cells seeded the previous week with wt-SS14 cells were trypsinized to allow the release and enumeration of spirochetes using the DFM. A total of 2-3×10^8^ treponemes were inoculated 24 hours after plating the Sf1Ep. Following treponemal addition, the total volume of media in each well was brought to 2.5 ml. Two days following treponemal cell addition, the plate was removed from the MA incubator, and 1 ml of old media was replaced with fresh one. Four days after treponemal cell inoculation, the plate was removed again from the MA incubator and the culture media was eliminated gently not to disturb Sf1Ep cells and adherent treponemes and replaced by 500 µl of transformation buffer (50 mM CaCl_2_, 10 mM Tris pH 7.4; equilibrated in MA) containing 15 ug total of p*tprA*arms-47p-*kan*^R^. As a control, to rule out CaCl_2_ toxicity to treponema cells, treponemes were also incubated with transformation buffer without plasmid vector. Cells were incubated in these transformation buffers (with and without plasmid) for 10 min at 34°C in the MA incubator and then washed twice with equilibrated TpCM-2 media to remove free plasmid from the culture wells. Finally, 2.5 ml of fresh TpCM-2 equilibrated in MA were added to the wells, and plates were returned to the MA incubator. The following day, concentrated tissue-culture grade liquid kanamycin sulfate (Sigma-Aldrich, St. Louis, MO) was added to the appropriate wells to reach a final concentration of 25 µg/ml. As a control, to confirm the treponemicidal activity of kanamycin, wild-type treponemes were also incubated in fresh TpCM-2 media containing 25 µg/ml of kanamycin sulfate. Kanamycin sulfate-containing TpCM-2 media was exchanged weekly but treponemes were sub-cultured every two weeks as per published protocol [11] until they reached a density of ∼3×10^7^ cells/ml, counted using the DFM, at which point they were sub-cultured first into one well of a 6-well plate (seeded with 10^5^ Sf1Ep cells on the previous day) to upscale the culture, and then into all the wells of a 6-well plate at the following passage to further expand the strain and minimize the chances of culture loss due to contamination. Whenever possible, treponemes that were not used for inoculation of a new plate were pelleted by centrifugation at 15,000 rpm for 10 min using a tabletop centrifuge and resuspended in 1X DNA lysis buffer (10 mM Tris-HCl, 0.1 M EDTA, 0.5% SDS), Trizol (Thermo Fisher), or used to make glycerol stocks, regardless of the number of treponemes counted by DFM.

To compare susceptibility to kanamycin of the *tprA*^ko^-SS14 and wild-type strains, treponemes obtained from the tenth *in vitro* passage were sub-cultured into two 96-well cell culture plates (Corning) instead of 6-well plates to allow a total of eight replicates for each kanamycin sulfate concentration tested. Briefly, the day before inoculation, two 96-well plates were seeded with 3×10^3^ rabbit Sf1Ep cells per well in 150 μl of culture media. The plates were then incubated overnight in the HeraCell incubator. On the same day, TpCM-2 media was prepared according to protocol and equilibrated overnight at 34°C in the MA incubator. On the following day, cell culture media was removed from the 96-well plates, and cells were rinsed with equilibrated TpCM-2 media. Subsequently, each well was filled with 150 µl of equilibrated TpCM-2 media, and the plate was transferred to the MA incubator. To prepare the treponemal inoculum for the 96-well plates, the Sf1Ep cells seeded the previous week with either the *tprA*^ko^-SS14 and wild-type strains were trypsinized to allow the release and enumeration of spirochetes. Treponemes were counted using the DFM and diluted in TpCM-2 to 3.3×10^5^ *T. pallidum* cells/ml to obtain a treponemal inoculum of 5×10^4^ cells in a total of 150 μl, which were then added to each well of the 96-well plates. The kanamycin sulfate concentrations tested were a 1:2 dilution series ranging from 200 to 1.6 μg/ml, for a total of 8 different concentrations tested in eight replicate wells. No-antibiotic wells, as well as solvent-only wells (water), were also included as controls. Tissue-culture grade kanamycin sulfate was purchased from Sigma-Aldrich. Wild-type and *tprA*^ko^-SS14 treponemes in no-antibiotic wells were harvested at day 0 (inoculum), day 1, day 4, and day 7 post-inoculation after incubation at 34°C in the MA incubator to assess treponemal growth in normal conditions and that, even in absence of antibiotic pressure, the *kan*^R^ gene would remain steadily integrated in place of the *tprA* pseudogene in the *tprA*^ko^-SS14 treponemes. The experiment where *tprA*^ko^-SS14 treponemes were grown in a 96-well plate in the presence or absence of kanamycin sulfate was performed twice to ensure a) the reproducibility, and b) to obtain DNA-free RNA to compare the level of transcription of the *kan*^R^ gene and that of the *tp574* genes by RT-ddPCR.

Treponemes harvested at passage #15 (Week 20 post-transformation) were resuspended in SDS-PAGE sample buffer and proteins were separated on a 12% pre-made acrylamide gel (Thermo Fisher) to evaluate the expression of the Kan^R^ protein using mass spectrometry after gel band excision and digestion (see protocol below).

### Rabbit infection, treatment, and sample collection

Pharmaceutical grade kanamycin sulfate (50 mg/ml in water) was prepared by Kelley-Ross compounding pharmacy in Seattle, WA, and stored at -20°C until use according to the pharmacist’s instructions. Once thawed, the bottle was kept at 4°C and removed from the fridge only to withdraw doses. On the day of infection, *tprA*^ko^-SS14 and wild-type strains were harvested from the wells of a 6-well culture plate, enumerated using a Nikon NiU darkfield microscope (Nikon, Melville, NY), and diluted in sterile saline to 3×10^7^/ml. Three NZW rabbits were infected. One rabbit was infected IT only in its left testicle with the *tprA*^ko^-SS14 strain. This rabbit received the first subcutaneous dose of kanamycin sulfate (5.0 mg/Kg) one hour before infection and was treated every 12 hours for a total of 10 days, each time with the same dose. The second rabbit was infected with the wt-SS14 strain and also treated with kanamycin as above, while the third rabbit was infected with wt-SS14 but not treated. When orchitis developed, rabbits were euthanized, and the left testicle was removed and minced in sterile saline to extract treponemes. The control rabbit infected with the wt-SS14 strain and treated with kanamycin was euthanized when the other control rabbits developed orchitis, and testicular extracts were also processed. Treponemal suspensions were spun at 1,000 rpm at 4°C in an Eppendorf 5430R refrigerated centrifuge, and cellular debris were discarded. Treponemes in the supernate were enumerated by DFM and resuspended in 1X DNA lysis buffer by mixing equal volume of extract and 2X buffer. The remaining treponemes were frozen in glycerol stocks (50% serum-saline + 50% sterile glycerol). Terminal bleeding through cardiac puncture was performed to obtain serum for VDRL and TPPA tests, performed as described above. All sera were heat-inactivated at 56°C for 30 min before use.

### DNA and RNA extraction

DNA extraction from cultured *tprA*^ko^-SS14 or wild-type strains propagated in 6-well plates following the transformation procedure was performed using the QIAamp mini kit (Qiagen) according to the manufacturer’s instructions. Extracted DNA was stored at -80°C until use for qualitative PCR, ddPCR, or WGS (see below). DNA extraction from the 96-well culture plates used to assess susceptibility to kanamycin of the *tprA*^ko^-SS14 and wild-type strain, respectively, was performed after a one-week incubation of the cells in the MA incubator. Plate culture media was removed with a vacuum manifold from each well and discarded. Cells were not trypsinized but 200 μl of Genomic Lysis Buffer (Zymo Research, Irvine, CA) for DNA extraction was added. Cells were then lysed through incubation in lysis buffer for 30 min at room temperature as per provided protocol. While propagating the SS14 strain, we used dark-field enumeration of treponemes present in the cell culture supernate and attached to Sf1Ep cells but released by cell trypsinization, and determined that the majority (∼85%) of *T. pallidum* cells *in vitro* adhere to the rabbit epithelial cell monolayer, similar to what was reported by Edmondson *et al.* for other cultivated strains [11]. This evidence allowed us to discard the culture media without concern that the experimental results would be significantly affected. Following cell lysis, the plates were frozen at -20°C until extraction could be completed. To purify DNA, the 96-well plates were thawed at 56°C in a dry incubator and quickly spun to recuperate condensation drops on the well lids. DNA was extracted using a Quick-DNA 96 kit (Zymo Research) according to the manufacturer’s protocol. DNA was eluted in 100 µl of water and stored at -20°C until amplification using qPCR to evaluate treponemal burden. The same samples were used to perform ddPCR (see protocol below) targeting *tp0001* (*dnaA*), *tprA*, and the *kan*^R^ genes to investigate the ratio between these targets. As an additional control, extracted samples were also tested randomly using pUC57-specific primers (Table 1) annealing upstream and downstream of the cloned insert to assess carry-over of the plasmid used for transformation.

RNA extraction was performed using the Quick-RNA 96 kit (Zymo Research) following the manufacturer’s instructions with the exception that in-column DNA digestion using the DNaseI enzyme (provided by the kit) was prolonged for a total of 1 hour and performed using 50% more enzyme than normally suggested. Single samples from *in vivo* and *in vitro* propagation were instead extracted according to the Trizol reagent manual. Total RNA was treated with DNaseI according to the protocol provided with the TURBO DNA-free kit (Thermo Fisher). DNA-free RNA was checked for residual DNA contamination by qualitative amplification using primers specific for the *tp0574* gene (primers in Table 1) as already described [44]. Reverse transcription (RT) of total RNA was performed using the High-Capacity cDNA Reverse Transcription kit (Thermo Fisher) with random hexamers according to the provided protocol. cDNA samples were stored at -80°C until use for qualitative PCR to demonstrate expression of the *kan*^R^ gene or for ddPCR to quantify the level of expression of the *tp0574* and the *kan*^R^ genes (primers in Table 1).

### Qualitative and quantitative PCR

Samples harvested during routine propagation of the *tprA*^ko^-SS14 and wild-type strains were assessed for integration of the *kan*^R^ gene into the *tprA* locus by using qualitative PCR. In the first amplification, the sense primer targeted a region of the *T. pallidum* genome immediately upstream of the left *tprA* homology arm of the vector (and hence not cloned into p*tprA*arms-*tp0574*prom-*kan^R^*), while the antisense primers targeted the *kan*^R^ gene, with the rationale that only a *kan*^R^ gene integrated into the *tprA* locus would provide amplification. Primers and amplicon size are reported in Table 1 and schematically represented in Fig.2, along with the results of the PCR done on DNA extracted from treponemes at Passage #7. In the second PCR, the sense primer targeted the *kan*^R^ gene, and the antisense primer (Table 1) targeted the genomic region downstream of the left *tprA* homology arm of the vector. Amplification of the *tp0574* gene was used as positive amplification control. Amplifications were performed using five microliters of extracted DNA in 50 μl final volume containing 2.5 units of GoTaq polymerase (Promega, Madison, WI), 200 μM of each dNTP, 1.5 mM of MgCl_2_, and 400 nM of sense and antisense primers. Cycling parameters were initial denaturation (94°C) and final extension (72°C) for 10 min each. Denaturation (94°C) and annealing (60°C) steps were carried on for 1 min each, while the extension step (72°C) was carried out for 1 or 2 min depending on amplicon length. A total of 40 cycles were performed in each amplification. A qPCR assay was instead used to quantify treponemal burden in samples extracted from the 96-well plate following the kanamycin susceptibility assay of the *tprA*^ko^-SS14 and wild-type strains. In this case, the treponemal burden was evaluated using a qPCR approach targeting the *tp0574* gene previously described [16]. Primers are reported in Table 1. Briefly, an absolute quantification protocol using an external standard was used to quantify the *tp0574* gene copy number at the time of sample harvest. Standard construction was also previously described in detail [16]. For amplification, the Powerup SYBR Green Master Mix (Thermo Fisher) was used. Amplifications were run on a QuantStudio 5 thermal cycler (Thermo Fisher) and results were analyzed using the instrument software. Data were imported into Prism 8 (GraphPad Software, San Diego, CA) and further analyzed to assess statistical significance of the values from test and no-antibiotic control groups using one-way ANOVA with the Dunnett test for correction of multiple comparisons or t-test, with significance set at *p*<0.05 in both cases.

### Droplet digital PCR (ddPCR)

Droplet digital PCR assays were conducted to assess the ratio between the number of copies of the *kan*^R^ gene and another target on *T. pallidum* genome in the *tprA*^ko^-SS14 strain, in this case, the *tp0574* gene, the *tp0001* gene (*dnaA*), or the *tprA* gene. The *tp0574* and *dnaA* targets were chosen because of their relative distance to the *kan*^R^ insertion site (< 10 Kbp from *dnaA*, and ∼513Kbp from the *tp0574* gene) to account for possible discrepancies due to the vicinity of the *kan^R^* gene to *T. pallidum* chromosomal origin of replication. ddPCR was also used to evaluate the level of transcription of the *kan*^R^ gene and the *tp0574* gene, as both genes are transcribed by the same promoter. Amplification of the *tprA* gene was used as control. To this end, four sets of primers/probes (Table1) were designed. DNA obtained from the 96-well plate used to perform the kanamycin susceptibility assay with the *tprA*^ko^-SS14 strain was used along with the DNA from the *tprA*^ko^-SS14 strain obtained from Passage #8, and DNA extracted from the *tprA*^ko^-SS14 and SS14 wild-type treponemes propagated *in vivo*. cDNA was obtained (as described above) from the 96-well plate used to perform the kanamycin susceptibility assay with the *tprA*^ko^-SS14 strain, and from the *tprA*^ko^-SS14 and SS14 wild-type treponemes propagated *in vivo* and *in vitro* (Passage #8).

ddPCR was performed on a Bio-Rad QX100 system (Bio-Rad, Carlsbad, CA). Each reaction was performed using ddPCR Supermix for Probes (Bio-Rad) with the final concentration of primers at 900 nM and probes at 250 nM in a total reaction volume of 25 µl. Before amplification, template DNA was digested with 25 units of EcoRI (New England Biolabs, Ipswich, MA). cDNA was used without the digestion step. After droplet generation, droplets were transferred to a 96-well PCR plate and amplified on a 2720 Thermal Cycler (Thermo Fisher) with the following cycling parameters: 94°C for 10 min, followed by 40 cycles of 94°C for 30 s and 60°C for 1 min, and 98°C hold for 10 min. After amplification, the plate was transferred to QX200 droplet reader (Bio-Rad). Results were analyzed using the QuantaSoft software (Bio-Rad).

### Analysis of Kan^R^ expression by mass spectrometry

Expression of the KanR protein (molecular weight of 31.1 KDa) was assessed by liquid chromatography-mass spectrometry (LC-MS) in *tprA*^ko^-SS14 and wild-type treponemes. For the SDS PAGE, the *tprA*^ko^-SS14 and wild-type treponemes were harvested as described above from culture plates (Passage 15, Week 20 post-transformation), counted by DFM, and centrifuged at 100xg (1,000 rpm in a tabletop centrifuge with a 9 cm radius) to remove residual Sf1Ep cells. The resulting supernate was then spun at 15,000 rpm at RT for 10 min to pellet treponemes and pelletresuspend in 1X SDS-PAGE sample buffer (50 mM Tris-HCl; 100 mM DTT; 70 mM SDS; 1.5 mM Bromophenol blue, 2M glycerol). Approximately 10^9^ treponemes were resuspended in a final volume of 200 µl of SDS-PAGE sample buffer. Samples were boiled and loaded onto a 12% precast Tris-Tricine gel in a mini-Protean apparatus (both from Bio-Rad, Hercules, CA). Gels were stained using SimplyBlue SafeStain (Thermo Fisher). Subsequently, a gel segment encompassing protein sizes between 20-40 kDa (and thus including the ∼30 kDa Kan^R^ protein) was excised and bands were subjected to overnight in-gel trypsin digestion as previously described [16, 24]. The volume of digestion products was reduced to approximately 10 μl using a speed-vac. Peptides were analyzed by LC-MS at the Fred Hutchinson Cancer Research Center proteomics facility using an LTQ HP1100 mass spectrometer (Thermo Fisher), results were analyzed using the Proteome Discoverer software. Identified peptides were filtered to a false discovery rate (FDR) of <0.01 to ensure high confidence in the identified peptides.

### Whole-genome sequencing

Whole-genome sequencing was performed following DNA extraction of *tprA*^ko^-SS14 and wild-type treponemes cultured *in vitro* (Passage #9; Fig.1). Pre-capture libraries were prepared from up to 100 ng input DNA using the KAPA Hyperplus kit (Roche) and TruSeq adapters and barcoded primers (Illumina), following the manufacturer’s protocols, yielding an average fragment size longer than 500 bp. Hybrid capture of *T. pallidum* genomic DNA was performed overnight (>16 hours) using a custom IDT xGen panel designed against the reference genome NC_ 010741, following the manufacturer’s protocol. Short-read sequencing was performed on an Illumina MiSeq with 192 bp single-end reads, yielding at least 1e6 reads per sample. Reads were adapter and quality trimmed using Trimmomatic v0.39 [45] and assembled to the TPA reference genome NC_021508.1, or NC_021508.1 with the *tprA* gene replaced by the kanamycin cassette, using Bowtie2 [46], to an average coverage exceeding 175x across the genome for all samples. Manual confirmation of expected coverage and junctions was performed by visual inspection in Geneious Prime v2020.1.2 [47].

### Derivation of a *tprA*^ko^-SS14 pure isolate by limiting dilution and increased antibiotic selective pressure

Sf1Ep cells were seeded into the wells of a 96-well culture plate at a density of 3,000 cells/well in 150 µl of MEM and cultured overnight in a 37°C in the 5% CO_2_ incubator. The next morning after the cells had adhered to the plate the MEM was exchanged for 135 µl TpCM2 media which had been equilibrated overnight in a 34°C in the MA incubator. The cells were equilibrated in the MA incubator for at least 3 hr before proceeding to inoculation of the *tprA*^ko^-SS14 cells, harvested using trypsinization from a 6-well plate where the strain was routinely propagated. Following harvest, cell concentration was determined by DFM. For the serial dilutions, 3.3×10^4^ *tprA*^ko^-SS14 cells were inoculated into each of 8 wells of the prepared 96-well plate, mixed thoroughly, and then 15 µl of the diluted *T. pallidum* mixture moved to the next set of 8 wells. In this manner, successive 1:10 dilutions of *tprA*^ko^-SS14 were made on the plate until the mixture was diluted to 0.33 *tprA*^ko^-SS14 cells/well. Kanamycin was added to each well for a final concentration of 200ug/ml. The plate was grown for 2 weeks in the MA incubator, with a half-media change at one week. After 2 weeks, the supernatant was removed from the plate, the wells washed briefly with 20ul of trypsin, and then incubated with 20ul of trypsin for 5min at 37°C to release adherent *T. pallidum* cells. Ten microliters of the resulting *T. pallidum* mixture were inoculated into a fresh 96-well plate prepared and incubated as described above. To the remaining 10 µl, 200 µl of genomic lysis buffer from a Quick-DNA 96 kit (Zymo Research, Irvine, CA) was added. The plate was sealed and stored at -20°C until DNA extraction. DNA was extracted using the Quick-DNA 96 kit (Zymo Research). DNA was eluted in 30 µl of molecular H_2_O, the minimum volume recommended for the kit. Lack of amplification signal for *tprA* was assessed by ddPCR as described above.

## Acknowledgments

This work was supported by the National Institute for Allergy and Infectious Diseases of the National Institutes of Health grant number U19AI144133 (Project 2 and Genomics and Isolation Core; Project 2 leader: L.G.; Core leaders: L.G. and A.G.; PI: Anna Wald, University of Washington). We are grateful to Dr. Philip Stewart, Ph.D. (NIH Rocky Mountain Laboratory, Hamilton, MT) and Mark C. Fernandez, Ph.D. candidate (University of Washington, Department of Global Health) for helpful suggestions on the experimental design. The senior author, Lorenzo Giacani, wishes to dedicate this milestone achievement in the field to Dr. Arturo Centurion-Lara, MD, mentor and friend, who prematurely passed away in 2018. The content of this study is solely the responsibility of the authors and does not necessarily represent the official views of the National Institutes of Health. The funders had no role in study design, data collection, and analysis, decision to publish, or preparation of the manuscript.

